# Inhibitory control of gait initiation in humans: an electroencephalography study

**DOI:** 10.1101/2024.01.25.577273

**Authors:** D. Ziri, L. Hugueville, C. Olivier, P. Boulinguez, H. Gunasekaran, B. Lau, M-L. Welter, N. George

## Abstract

Response inhibition is a crucial component of executive control. Although mainly studied in upper limb tasks, it is fully implicated in gait initiation. Here, we assessed the influence of proactive and reactive inhibitory control during gait initiation in healthy adult participants. For this purpose, we measured kinematics and electroencephalography (EEG) activity (event-related potential [ERP] and time-frequency data) during a modified Go/NoGo gait initiation task in 23 healthy adults. The task comprised Go-certain, Go-uncertain, and NoGo conditions. Each trial included preparatory and imperative stimuli. Our results showed that go-uncertainty resulted in delayed reaction time (RT), without any difference for the other parameters of gait initiation. Proactive inhibition, i.e. Go uncertain versus Go certain conditions, influenced EEG activity as soon as the preparatory stimulus. Moreover, both proactive and reactive inhibition influenced the amplitude of the ERPs (central P1, occipito-parietal N1, and N2/P3) and theta and alpha/low beta band activities in response to the imperative—Go-uncertain versus Go-certain and NoGo versus Go-uncertain—stimuli. These findings demonstrate that the uncertainty context induced proactive inhibition, as reflected in delayed gait initiation. Proactive and reactive inhibition elicited extended and overlapping modulations of ERP and time-frequency activities. This study shows protracted influence of inhibitory control in gait initiation.

## 1. INTRODUCTION

The adaptation of motor behaviour to environmental constraints is essential for performance and survival. Depending on the situations encountered, this implicates to stop or refrain upcoming behaviour, which involves response inhibition processes. Two different, context-dependent mechanisms of inhibition have been described. The first one occurs before any action is initiated and is called “proactive inhibition” (Albares et al., 2015; Criaud et al., 2012). The movement can be initiated only once this mechanism is released. Proactive inhibition is activated in uncertain contexts. By delaying any movement, it allows time for deciding on and selecting the appropriate response to initiate. The second mechanism takes place when the action to be initiated is prepared and is called “reactive inhibition” (Aron et al., 2007; Chambers et al., 2009; Verbruggen & Logan, 2008). It allows stopping the movement when a STOP signal is perceived. Behavioural, neurophysiological, and neuroimaging studies in healthy adults have characterized inhibitory processes (Albares et al., 2014; Boulinguez et al., 2009; van Belle et al., 2014, for review see: Bari & Robbins, 2013) and their modulation in different cognitive and emotional contexts (Pessoa et al., 2012; Shafritz et al., 2006). However, they have mainly used tasks including a mixture of Go and NoGo or Stop stimuli, thus concentrating on reactive inhibition, with simple motor responses of the upper limb, such as button presses. Such tasks have the advantage of being simpler to use in the laboratory settings, yet they may not give access to the full dynamics of inhibitory processes and their interplay with online motor control (Hervault et al., 2022), which is crucial for more complex behaviour, such as gait initiation. Here, we were interested in characterizing proactive and reactive inhibitory processes during a gait initiation task in healthy middle-aged adults. This will be key for future endeavour to understand gait impairment in neurodegenerative diseases such as Parkinson’s disease.

It has been proposed that proactive inhibition may constitute a “default mode” of the brain (Criaud et al., 2012). It would encompass two types of processes: pre-amping, that is, amping up reactive inhibition in preparation to possible upcoming Stop or NoGo stimuli, and pre-setting, that is, presetting action control to enhance reactive inhibitory success in response to Stop and NoGo stimuli (van den Wildenberg et al., 2022). Both processes may be active under the classical conditions of Go/NoGo and Stop Signal tasks and specific paradigms are therefore required to uncover the effects of proactive inhibition. Accordingly, Albares et al. (2014) proposed a modified Go/NoGo task including a “Go-certain” condition wherein subjects had to systematically press a button in response to a visual stimulus, in addition to the “Go-uncertain” condition where both Go and NoGo trials were included and intermixed. They showed that the reaction time was increased in the context of Go-uncertain trials as compared to Go-certain trials (Albares et al., 2014; see also Criaud et al., 2016). This delay was considered to reflect the time needed for lifting proactive inhibition in the Go-uncertain condition. It may be associated with the action pre-setting process of proactive inhibition in this Go-uncertain condition relative to the Go-certain one (van den Wildenberg et al., 2022).

The most known electrophysiological signature of inhibition is the event-related potential (ERP) N2/P3 complex, observed in electroencephalography (EEG) studies. In Go/NoGo tasks of the upper limb, the comparison between Go and NoGo trials revealed a fronto-central negativity peaking at 200-400 ms after the stimulus (N2), which is larger (aka. more negative) when inhibition is required (Eimer, 1993; S. Kaiser et al., 2006; Pfefferbaum et al., 1985). It is followed by a positivity at 300-500 ms (P3), which typically peaks on frontal electrodes in response to NoGo stimuli, whereas it has a centro-parietal peak in response to Go stimuli (Falkenstein et al., 1999). This distinction is akin to the P3a / P3b distinction; NoGo P3 is considered a P3a-like component, while the centro-parietal P3 for Go stimuli is akin to P3b in response to targets in oddball paradigms. These components have been proposed to reflect different executive, neuroinhibitory mechanisms, with P3a considered as reflecting the involuntary relocation of attention and P3b the process of cognitive control (for reviews, Pires et al., 2014; Polich, 2007). The NoGo P3 is also elicited after the stop stimulus during Stop Signal tasks (Waller et al., 2021; Wessel, 2018). It is considered as a landmark of reactive inhibition (for review see Polich, 2007). Moreover, in the frequency domain, modulations of theta power (Harper et al., 2014; Messel et al., 2021), alpha power (Albares et al., 2014) and beta power (DeLaRosa et al., 2020; Picazio et al., 2014) over frontal and central regions have been reported during Go/NoGo and Stop Signal tasks. While these activities were initially considered mainly in association with reactive inhibition, they are also modulated in protocols emphasizing proactive inhibition (Albares et al., 2014; Messel et al., 2021; Pscherer et al., 2023; Soh et al., 2021). These activities were associated with activations of the pre-supplementary motor area (pre-SMA), the inferior frontal cortex, and the basal ganglia (for review, see Aron, 2011; Swann et al., 2009; Swann et al., 2012). Furthermore, comparing Go-certain and -uncertain conditions has allowed identifying a fronto-central component of the ERP response, which was associated with proactive inhibition. This so-called dMF170 component was maximum around 170 ms and its amplitude on the fronto-central electrode (FCz) where it peaked was inversely correlated with the reaction time to Go stimuli (Albares et al., 2014). The source of dMF170 was localized to the pre-SMA, in agreement with the fMRI studies that associated proactive inhibition with activation of the pre-SMA and SMA (Albares et al., 2014; Criaud et al., 2016).

All the above-mentioned studies were based on upper-limb tasks. In daily life activity, walking in various environment represents a highly challenging situation where locomotion should be initiated (Go), or stopped (NoGo), in various contexts putting different demands on proactive and reactive inhibitory control processes. For example, crossing a street is likely to involve proactive inhibition while avoiding the collision with a scooter circulating on the walkway involves reactive inhibition. Moreover, successfully initiating gait depends on specific anticipatory postural adjustment (APAs) that allow discharging the swinging leg from body weight in order to lift the first foot, while maintaining postural stability to avoid falling (Breniere & Do, 1986; Jian et al., 1993). This gait initiation process is controlled, at least partly, by a brain network including the SMA and pre-SMA cortices, basal ganglia and mesencephalic locomotor region (Gilat et al., 2017; Lau et al., 2019; Marchal et al., 2019; Masdeu et al., 1994; Nadeau, 2007; Varghese et al., 2016). It can be affected in various pathological conditions such as Parkinson’ disease, where gait initiation alteration can lead to freezing of gait (FOG), an inability to lift the foot from the floor, which worsens when the patient is placed in a cognitively or emotionally demanding situation (Beaulne-Séguin & Nantel, 2016; Lagravinese et al., 2018; Spildooren et al., 2010). The fact that FOG mainly occurs at gait initiation could reflect increased inhibitory processes with an inability to release the motor program to start walking. This pathological phenomenon thus reflects the tight interaction between inhibitory and motor processes and the importance of understanding their interplay during tasks such as gait initiation. In particular, it is unclear if inhibitory mechanisms uncovered for upper limb movements, such as simple finger movements, may generalise to gait initiation, considering that the associated motor circuits are partly different and several studies suggest that this generalisation is not trivial. On one hand, global or non-selective inhibition of movements may involve some common neural mechanisms (Goode et al., 2019), on the other hand, there may be inhibition processes specific of the movement to be inhibited (Hervault et al., 2019, 2021, 2022; see Hannah & Aron, 2021 for review).

Here, we studied gait initiation in healthy adult humans using a modified Go/NoGo task derived from Albares et al. (2014). Our primary aim was to disentangle proactive and reactive inhibition processes and characterizing their neurophysiological signatures with EEG. Our hypothesis was that gait initiation, as reflected by the onset of APAs, would be delayed in the Go-uncertain relative to the Go-certain conditions. We also explored the potential impact of Go-uncertainty on other parameters of gait initiation, such as APA duration and amplitude, as well as the reaction time for the first foot-off. For EEG activities, given that the interplay of inhibitory mechanisms and motor control during gait initiation was unknown, we used a data driven approach, followed by analysis of the ERPs and time-frequency activities in time windows of interest. Our study is an initial stride towards understanding how proactive and reactive inhibitory processes influence gait initiation in healthy adult human subjects.

## 2. MATERIAL AND METHODS

### 2.1. Participants

Twenty-five healthy adult subjects (10 women / 15 male) participated in this study. This sample size was based on Albares et al. (2014) (n=20) and considering resource constraints. The inclusion criteria were to be healthy adults, aged between 35 and 70 years, without any previous or ongoing, orthopaedic or neurological medical history. The subjects were recruited from the healthy volunteers list of the Clinical Investigation Centre located at the Paris Brain Institute. Two participants were excluded from analysis due to technical issues, leading to a final sample of 23 subjects (10 women; mean age: 52 ± 15 years). All subjects provided written informed consent and received financial compensation for their participation. This study was performed according to the Declaration of Helsinki and good clinical practice guidelines; it was legally sponsored by INSERM (RBM nb. C11-40; N°IDRCB 2012-A00225-38) and approved by a local ethics committee (CPP Paris VI, project nb. 20-12; ClinicalTrials.gov ID: NCT01682668).

### 2.2. Experimental protocol

In this study, we assessed whole-head EEG activity and kinetics parameters of gait initiation during a modified Go/NoGo task.

For this purpose, we used a task previously validated with upper limb movements (Albares et al., 2014). In this task, the subject had to initiate (Go condition) or to refrain from initiating gait (NoGo condition) in response to a visual stimulus (Figure 1). There were two contexts of Go condition presented in separate blocks:

- In the Go-certain condition, only Go trials were presented. The trials started with a green plus sign preparatory cue presented for 100 ms, followed by the imperative Go stimulus (a filled green circle, presented for 100 ms) after a random interstimulus interval (ISI) of 1, 1.5 or 2 s
- In the Go-uncertain condition, both Go and NoGo trials were presented. The trials started with a red plus sign followed by either the green filled circle Go stimulus or a green cross sign, which was the NoGo stimulus. The timing of stimulus presentation was the same as in the Go-certain condition. The Go-uncertain and NoGo trials were in 50/50 proportion and presented in a pseudo-randomized order, with no more than 3 consecutive identical trials.

**Figure 1.**
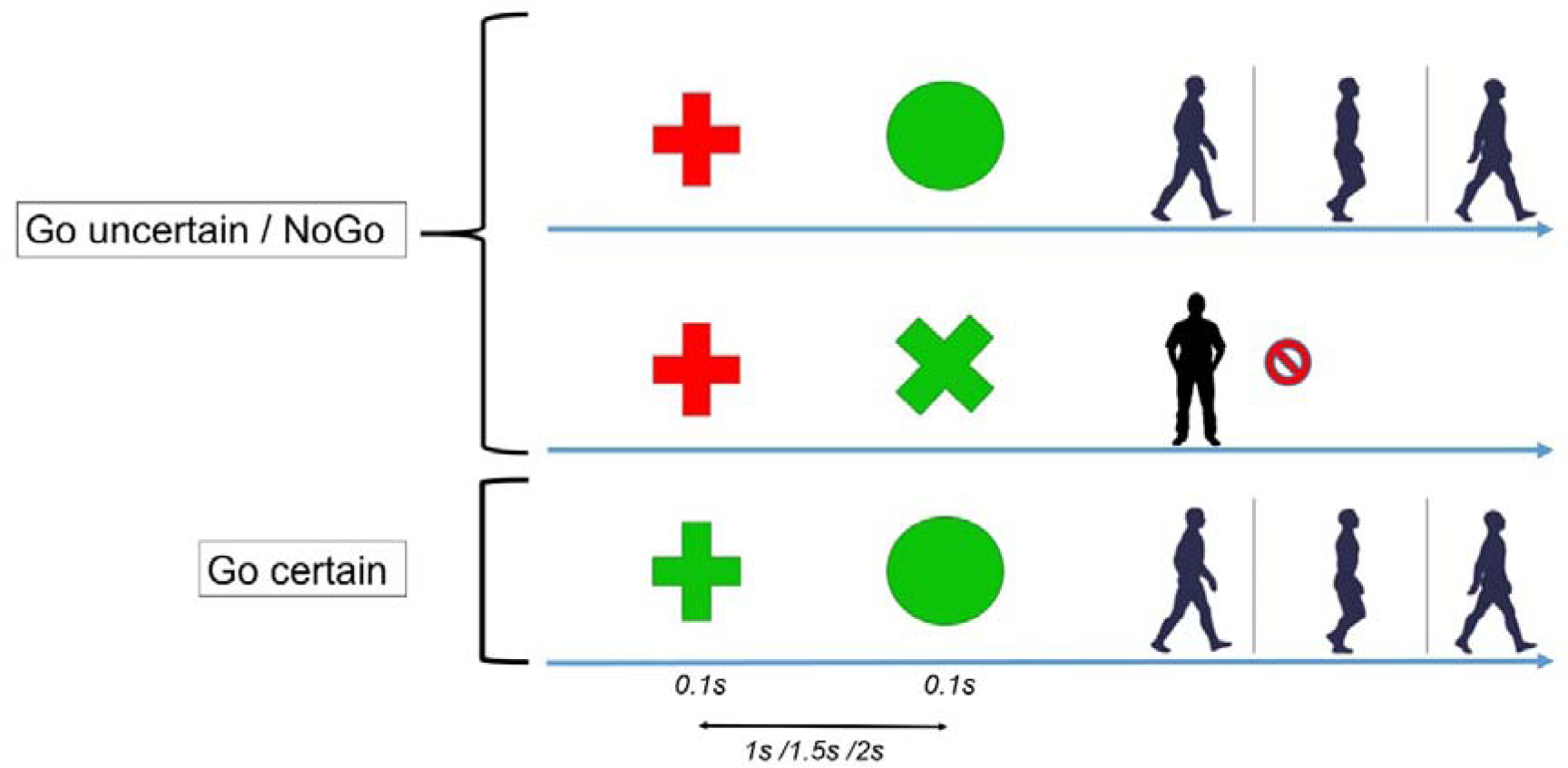
Experimental design. Illustration of the modified Go/NoGo task including a Go-certain condition, which was used to disentangle proactive and reactive inhibition.

For both Go-certain and -uncertain conditions, the subject had to initiate walking at their self-paced gait speed as soon as they perceive the Go stimulus. Conversely, they were instructed to refrain from any movement and to maintain a stationary posture upon perceiving the NoGo stimulus. The instructions insisted on not anticipating movement and refraining from initiating gait before the Go stimulus presentation. During the Go trials, the subject walked approximately 4 meters, executed a U-turn, and returned to their initial position to await the subsequent trial. Each trial was triggered manually by the experimenter after the subject had returned to the starting point and stabilized their stance, ensuring minimal displacement of the center of pressure (CoP), which was continuously monitored and visualised in real-time. The visual stimuli (spanning a surface area of 50x50 cm^2^) were projected onto the center of a white wall positioned 4.75 m in front of the subject by a ceiling mounted video projector within the experimental room.

The experiment started with a block of 10 Go-certain trials, followed by a block of 20 Go-uncertain / 20 NoGo trials (in pseudo-randomized order) and then a block of 10 Go-certain trials. This sequence of three blocks was repeated twice, for a total of 60 Go-certain, 60 Go-uncertain, and 60 NoGo trials (except for one subject who performed only two blocks of trials).

### 2.3. Data acquisition

#### 2.3.1. Recording of gait initiation kinetics parameters

Gait initiation kinetics parameters were recorded using a force platform (0.9x1.8m, Advanced Mechanical Technology Inc. LG6-4-1, sampling frequency at 1kHz) and 13 infrared cameras (Vicon system, 1.3 megapixels, sampling frequency 200Hz). The subject was dressed in shorts, tank top, and barefoot with 35 reflective markers positioned on the trunk, pelvis, and four limb joints. They initiated gait on the force platform that recorded the moments and forces in the 3-axis and allowed the calculation of the centre of foot pressure displacements (CoP) and centre of mass (CoM) velocity. In this study, we analysed the displacements of the markers placed on the heels and the 1^st^ and 5^th^ toes. For this purpose, the recorded markers were automatically reconstructed and manually labelled through Vicon Nexus 2.10.3 software interface, and the heel-off and the foot-off events of each gait initiation trial were identified. We reviewed each trial and placed manually the gait initiation events using homemade a Matlab R2019b script that allowed aligning and visualising simultaneously the heel-off trajectory in a vertical axis and the antero-posterior (AP) and medio-lateral (ML) displacements of the CoP (Supplementary Figure 1). We rejected the trials missing due to software glitch and the trials without clear baseline regarding CoP.

We identified and marked three events: 1) the first displacement of the CoP on the medio-lateral axis (t0) corresponding to the onset of the anticipatory postural adjustments (APAs), 2) the foot-off (FO) and 3) the foot-contact (FC) of the first step. The following parameters were then calculated: 1) the APA reaction time (APA RT), corresponding to the delay between the Go signal and t0, 2) the APA phase duration, corresponding to the delay between t0 and FO, 3) the FO reaction time (FO RT), corresponding to the delay between the Go stimulus and the foot-off, 4) the amplitude of the AP and ML CoP displacements during APAs, and 5) the step length, corresponding to the anterior CoP displacement between t0 and FC, and step width, corresponding to the mediolateral CoP displacement between t0 and FC.

#### 2.3.2. EEG data acquisition and analysis

EEG was recorded using an Easycap net with 128 active electrodes (actiCAP, Brain Products GmbH, Germany) placed according to the international extended 10-5 system and connected to actiCHamp amplifiers (Brain Products GmbH, Germany). The EEG signal was continuously recorded throughout every block of the task with a sampling rate of 25kHz and a band pass filter of DC-6100Hz. Electrode FCz was used as the reference electrode, leaving a total of 127 recorded channels, and FPz was the ground electrode. We checked electrode impedance aiming at electrode impedance below 10 kOhms; signal quality was further checked visually.

EEG data analysis was performed using the Fieldtrip toolbox (Fieldtrip-20200831; Oostenveld et al., 2011) running in Matlab R2019b in Linux environment. Data were first down sampled at 500Hz and a notch filter at 50, 100 and 150 Hz was applied. The continuous EEG data were filtered between 1 and 40 Hz, and then epoched between -500ms before the start of the trial to +700 ms post Go/NoGo stimulus. This epoching allowed rejecting the periods grossly artefacted by gait. The epochs were reviewed using the summary and ft_databrower functions of Fieldtrip, to reject artefacted epochs and mark electrodes with noisy signals that could interfere with independent component analysis (ICA). We then performed ICA to identify visually and reject the ICs associated with eye blinks and – when identifiable – horizontal eye movements (mean number of rejected components per subject +/- SD = 2.26 +/- 0.9; range = 1 to 5 ICs out of 108 to 127 total ICs per subject and experimental block; this total number of ICs was equal to the total number of retained electrodes). For some subjects, there were some ICs related to muscle or to movement artefacts. These ICs were not removed, because muscle and movement-related artefacts tend to be not stationary, hence they are unsatisfactorily corrected by ICA, and they were not identified in every subject. Rather, we reviewed visually the corrected data to reject trials with remaining artefacts and the signals from bad electrodes were reconstructed by interpolation from nearest neighbors. The mean number [SD] of rejected trials by subject was 4.2 [2.6] in the Go-certain condition, 2.3 [2.0] in the Go-uncertain condition, and 2.5 [2.7] in the NoGo condition. The mean number [SD] of reconstructed channels per subject was 3.4 [3.8]. Finally, the data were re-referenced according to an average reference to have an approximation of the true voltages over the head (Picton et al., 2000) and to reconstruct the signal on FCz electrode (as the average signal of all other electrodes).

We performed event-related potential (ERP) analysis and time-frequency (TF) decomposition and analysis of the EEG signal. These analyses were performed in two time periods, centered respectively around the preparatory cue stimulus onset and the imperative (Go / NoGo) stimulus onset. Thus, we defined new epochs: 1) between 800 ms before and 1.3 sec after the preparatory cue onset for the analysis of cue-related activities and 2) between 500 ms before and 700 ms after the imperative stimulus onset for the analysis of Go / NoGo-related activities.

For ERP analysis, the data were low-pass filtered at 30 Hz and baseline-corrected using the 200 ms before stimulus onset (preparatory cue or imperative Go / NoGo) in each time window of analysis. ERP were averaged separately for each experimental condition (Go-certain, Go-uncertain, NoGo) for each subject. We also computed the grand average of the ERPs across the 23 subjects for figure purposes.

For TF analysis, the data in each time period of analysis (centered on the cue and on the Go / NoGo stimuli respectively) were decomposed in the time-frequency domain between 1 and 40 Hz using the multi-taper method (mtmconvol function of Fieldtrip). We used frequency bins of 1 Hz, TF computation windows that slide by steps of 50 ms, with a Hanning window and a frequency resolution of 3 cycles per window (leading to varying time windows of TF transform for different frequencies: time window of 1s at 3 Hz and of 100 ms at 30 Hz). The data were then averaged separately for each experimental condition (Go-certain, Go-uncertain, NoGo) and baseline-corrected using a log ratio, considering the mean of the TF data between -500 and -100 ms before the stimulus as the baseline for each time period of analysis (centered on the preparatory cue and on the imperative Go/NoGo stimuli respectively). The data averaged by condition but not baseline corrected were also saved for the purpose of the analysis on the pre-cue stimulus period (see below). We also computed the grand average of the TF data across the 23 subjects for figure purposes.

### 2.4. Statistical analysis

For gait initiation parameters, we calculated the data distribution (mean and standard deviation) and assessed the differences in gait initiation parameters (APA and FO RT, anteroposterior and mediolateral amplitude of the APAs, APA duration, length and width of the first step) between Go-certain and Go-uncertain conditions, across subjects. Initial Shapiro-Wilk tests on each parameter indicated that these data were not normally distributed. Thus, we used non-parametric Wilcoxon’s signed-rank tests for our comparisons. We used a Bonferroni correction for multiple comparisons, setting the level of significance at p < 0.0071 (that is, p < 0.05 / 7, corresponding to the number of analysed gait parameters).

For the EEG data (ERPs and TF data), we used data-driven cluster-based approaches, as implemented in FieldTrip (Maris & Oostenveld, 2007). Thus, for each time period (centered on cue and on Go/NoGo stimuli, respectively), for every channel/time sample (for ERPs) or every channel/time/frequency sample (for TF data), we compared our conditions by means of paired t-test whose alpha threshold was set at p<0.01. To exclude any remaining artefact that may bias the results, we excluded the lowest rows of electrodes from these analyses (excluded electrodes: Fp1, Fp2, AFp1, Afp2, AF7, AF8, F9, F10, P9, O9, PO9, Iz, O10, PO10, P10, TP9, TP10, TPP9h, PPO9h, OI1h, OI2h, POO10h, PPO10h, TPP10h). The minimum number of electrodes to form a cluster was set at 2 and clusters were formed based on temporal (for ERPs) or temporal and frequency (for TF data) adjacency. We used sum(t) as the cluster statistics. To establish the distribution of this cluster statistics under the null hypothesis, we then used the Monte-Carlo method with 1000 permutations, as follows: For each permutation, the labels of the conditions were randomly permuted on a within-subject basis; paired t-tests were performed as for the original data; the sum(t) of observed clusters were computed, and the maximal value of the sum(t) [max(sum(t))] was retained. This was repeated 1000 times. This allowed building the distribution of max(sum(t)) under the null hypothesis. The clusters obtained from the original data were then compared to this null distribution and considered as significant if their sum(t) value exceeded the 97.5^th^ percentile or did not surpass the 2.5^th^ percentile (aka. corrected p<0.05 in two-sided test) of the max(sum(t)) Monte -Carlo distribution and they lasted at least over two time points. These cluster-based analyses were performed: 1) for the comparison of the Go-uncertain versus Go-certain conditions between 0 and 500 ms post-cue and between 0 and 500 ms post-Go stimuli, for both ERPs and TF data. In addition, for TF data, we analysed the pre-stimulus period between -500 and 0 ms before the cue onset using the TF data without baseline correction; 2) for the comparison of the NoGo versus Go-uncertain conditions between 0 and 500 ms post-Go/NoGo stimuli, for both ERPs and TF data.

The cluster-based analyses revealed ERP and TF differences in response to Go-uncertain versus Go-certain stimuli and to NoGo versus Go-uncertain stimuli, which were widespread in time and electrodes. In order to further characterize these differences and facilitate their interpretation, we extracted the mean amplitude around the maximum of each ERP component (central P1, occipito-parietal N1 and P2, central N2, occipito-parietal and frontal P3) or frequency band (theta [3-7 Hz], alpha [8-12 Hz], beta [13-21 Hz]) encompassed in these widespread clusters (see Results for details) and plotted it across subjects in each condition. We performed complementary Student t-tests on these mean amplitudes to further confirm the differences shown by the cluster-based analysis. Note that these analyses were purely confirmatory and for illustration purpose, and they are not reported in the results to avoid double dipping.

## 3. RESULTS

### 3.1. Effect of uncertainty on gait initiation parameters

We recorded and analysed a total of 3444 gait trials, with a mean [SD] of 47 [10] analysed trials per subject for the Go-certain condition and 52 [10] trials per subject for the Go-uncertain condition. All subjects performed the gait task adeptly, with scarce errors: there were only 3 trials in total with omission errors (no gait initiation upon a Go stimulus) and 2 trials with commission errors (initiating gait with foot-off despite the NoGo stimulus). In addition, in response to the NoGo stimulus, we observed sometimes a small lateral shift of the CoP not followed by foot-off (see Supplementary Figure 1). These “pre-APAs” (Delval et al. 2012) were constitutive of partial errors and observed in a mean [SD] of 17.5 [14] trials per subject.

The APA RT was significantly shorter in the Go-certain (mean [SD] = 117 [123] ms) than the Go-uncertain conditions (174 [124] ms), with a mean difference [SD] of 57 [37] ms (standardized Wilcoxon’s test value, z = -4.79, p = 2.4.10^-6^) (Figure 2). Additionally, the FO RT was significantly shorter in the Go-certain (636 [151] ms) than the Go-uncertain conditions (720 [152] ms), with a mean difference [SD] of 84 [40] ms (z = -5.04, p-value = 4.8.10^-7^). We did not observe any other significant difference in gait initiation parameters (APAs duration, AP and ML CoP displacements during APAs and first step length) between the Go-certain and -uncertain conditions (Figure 2; Supplementary Figures 2 and 3).

**Figure 2.**
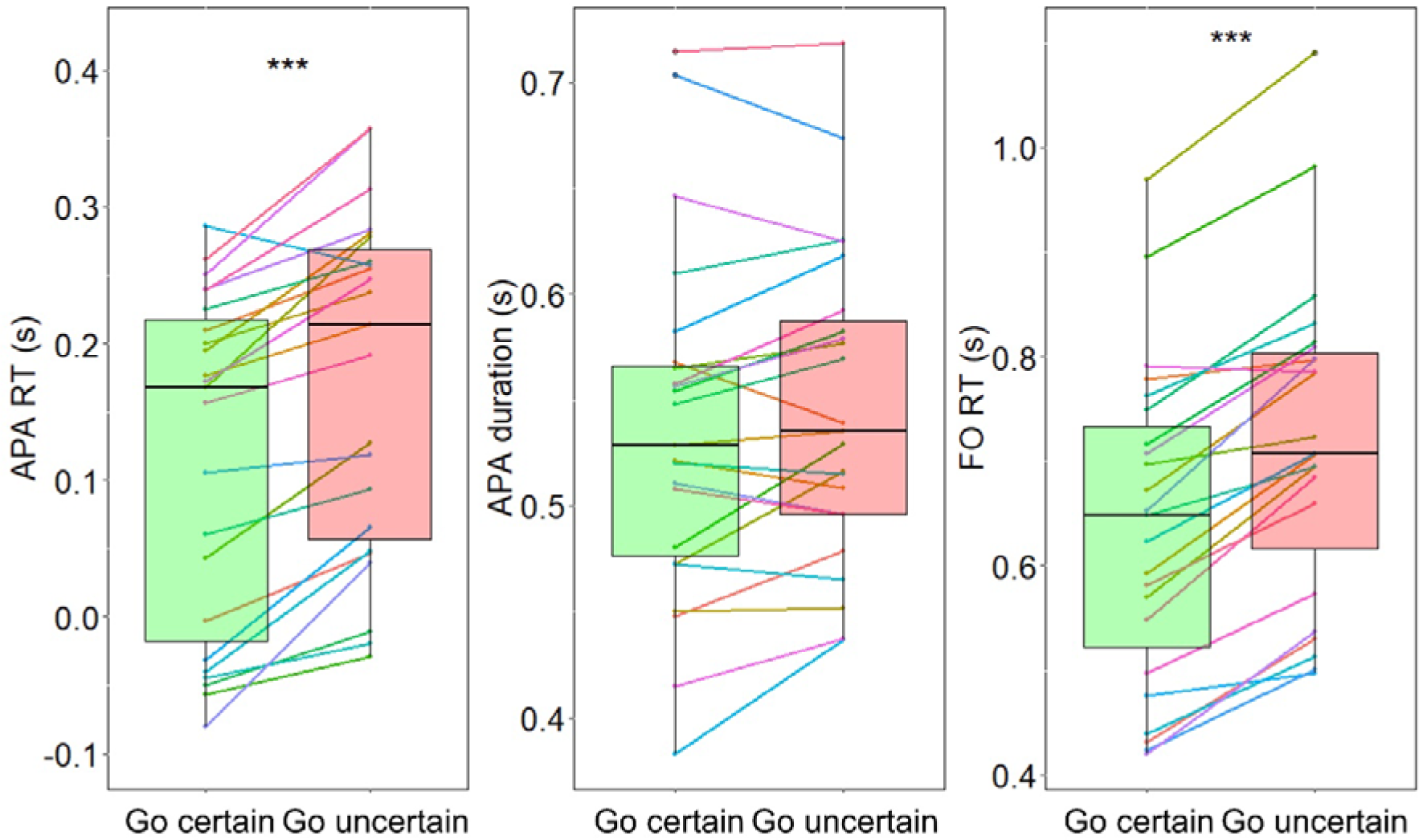
Reaction time during gait initiation in the Go-certain and Go-uncertain conditions. The main parameters of gait initiation for Go-certain (in green) and Go-uncertain (in red) trials are represented. From left to right: APA RT, that is, the delay from the Go stimulus to the start of the medio-lateral CoP displacement; APA duration; FO RT, that is, the delay from Go stimulus to the first foot-off. The boxplots encompass the second and third quartiles of the data, with the median shown as a thick black horizontal line; the vertical thin line represents the minimum and maximum values of the individual data. The colored lines between the Go-certain and Go-uncertain boxplots represent the individual data. The significant differences are indicated by asterisks (***: p<0.0001).

Upon examining individual gait parameters, we noted a mean APA-RT below 0 in the Go-certain trials in 7 out of the 23 subjects, with an anticipation of gait initiation (i.e. APA onset prior to imperative stimulus) in more than half of the trials in those subjects (average number of anticipated trials = 44.6). This also included a mediolateral displacement of the CoP following the preparatory cue in some trials. Three out of these 7 subjects also showed a negative mean APA-RT in the Go-uncertain trials (with an average number of anticipated Go-uncertain trials = 42.3). However, for all these “anticipated” trials, the FO occurred after the presentation of the Go stimulus. This indicated that although these subjects anticipated gait initiation, they refrained from lifting their foot from the ground until the appearance of the Go stimulus. When we excluded these “anticipated” trials from the analysis, the mean APA RT remained significantly lower in the Go-certain than the Go-uncertain trials (z = -4.46, p = 8.4.10^-6^).

### 3.2. Event-related potential analysis

We identified a sequence of well-established ERP components following both the preparatory cue and the imperative stimuli. First, the visual P1 peaked around 100 ms. It was accompanied by a focal central positivity (referred to as central P1). Second, a large occipito-parietal N1 peaked between 150 and 200 ms, followed by an occipito-parietal P2. Third, we observed the N2/P3 complex. The N2 component reached its maximum in fronto-central regions between 250 and 350 ms, and the P3 exhibited a peak in occipito-parietal and/or frontal regions, contingent upon the experimental condition, between 350 and 500 ms (Figures 3 and 4).

**Figure 3.**
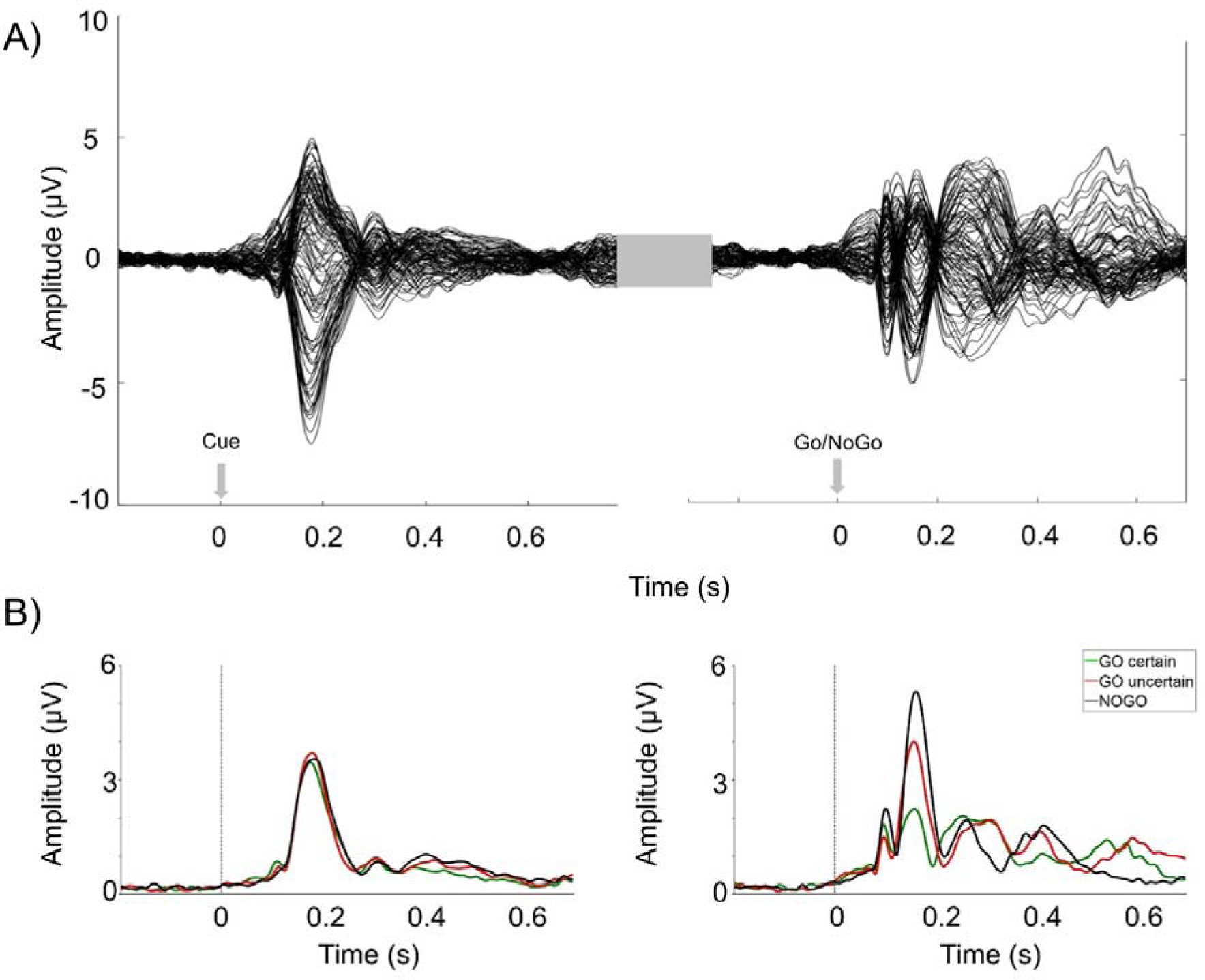
Overall time course of ERPs in response to the preparatory (Cue) and imperative (Go / NoGo) stimuli. A) The overall mean across subjects of the ERPs (in µV) across time (in s) is represented on all analysed electrodes overlaid. On the left, ERPs in response to the preparatory stimulus are represented. On the right, ERPs in response to the imperative stimulus are represented. These ERPs are here illustrated in the Go-certain condition. B) The amplitude (aka. root mean square, in µV) of the global field power across time (in s), in response to the preparatory stimulus (on the left) and to the imperative stimulus (on the right), is represented in the three conditions. The Go-certain condition is in green, the Go-uncertain in red and the NoGo in black.

**Figure 4.**
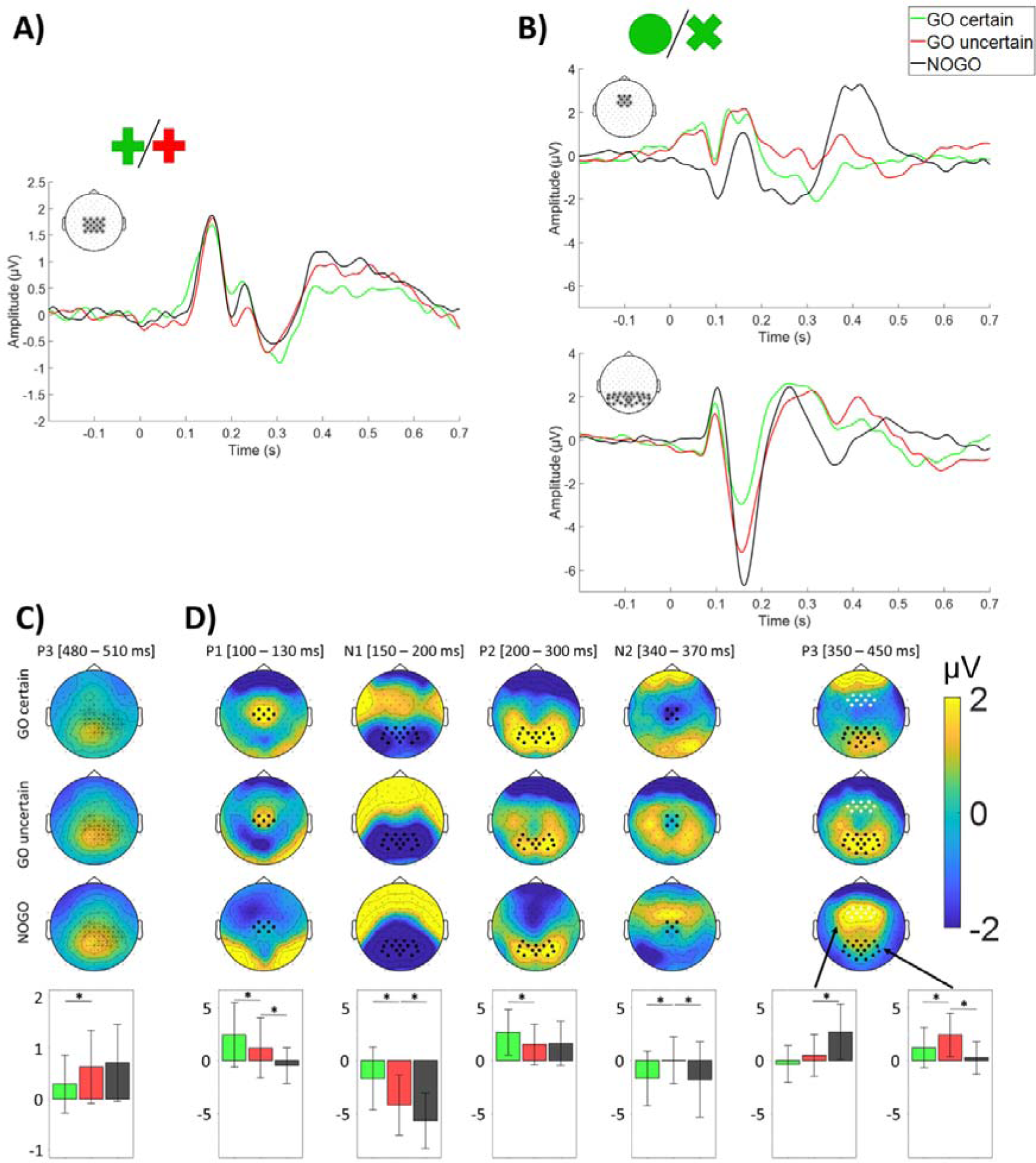
ERPs in response to the preparatory (Cue) and imperative (Go / NoGo) stimuli in the Go-certain, Go-uncertain and NoGo conditions. <u>Upper part:</u> Time course of ERPs in response to the preparatory cue stimulus (A) and to the imperative stimulus (B). The grand mean of the ERPs across subjects, averaged over the electrodes indicated by a ‘*’ on the topographical map inset is represented for the Go-certain (in green), Go-uncertain (in red) and NoGo (in black) conditions. <u>Lower part:</u> Topographical maps of the ERP components encompassed in the observed significant clusters (see text for details) and bar plots of the mean amplitude of these components, for the Go-certain, the Go-uncertain, and the NoGo conditions. C) The uncertain relative to the certain contexts were associated with a larger P3 in response to the preparatory stimulus. The electrodes marked by a ‘•’ are the cluster’s electrodes that were used to draw the bar plot underneath the map. In this bar plot, the overall mean amplitude of the P3 across subjects, on these electrodes and in the time window indicated in square brackets, is represented for the Go-certain (in green), the Go-uncertain (in red), and the NoGo (in black) conditions. The statistically significant difference between Go-certain and uncertain condition is indicated by a ‘*’. The thin vertical lines represent the standard deviation across subjects. D) There were modulations of the central P1, N1, P2, and N2/P3 complex in response to the imperative Go-certain, Go-uncertain, and NoGo stimuli. On each map, the electrodes used to draw the bar plots underneath the map are marked by a black ‘•’ for P1, N1, P2, N2, and parieto-occipital P3, and by a white ‘•’ for frontal P3. Below the maps, the grand mean amplitude of these components across subjects, on the selected electrodes and in the time window indicated in square brackets next to the component label, is represented as bar plots for the Go-certain (in green), the Go-uncertain (in red), and the NoGo (in black) conditions. The statistically significant differences between conditions are indicated by a ‘*’. The thin vertical lines represent the standard deviation across subjects.

#### 3.2.1. Effect of proactive inhibition: comparison between Go-certain and Go-uncertain trials

The cluster-based analysis revealed a single statistically significant cluster of difference between the two conditions during the cue period. The amplitude of the P3 was larger in response to the Go-uncertain preparatory cues compared to the Go-certain ones (mean difference [SD] = 0.34 [0.55] µV; Figure 4A and 4C). This was reflected by a significant centro-parietal cluster between 480 and 510 ms (sum(t)=-584; p=0.037; Supplementary Table 1 and Supplementary Figure 4).

In contrast, the ERPs in response to the imperative Go-certain and Go-uncertain stimuli exhibited sustained differences across time and electrodes, reflected by several large, significant, positive and negative, clusters (all corrected p_s_ < .05). For the sake of clarity, we describe these effects in relation to the modulated ERP components (Figures 4B and 4D). First, the early central and focal P1 was reduced for the Go-uncertain compared to the Go-certain trials, with a mean amplitude difference [SD] of 1.23 [0.94] µV on central electrodes between 100 and 130 ms. Second, the occipito-parietal N1 was larger (more negative) for the Go-uncertain than the Go-certain trials, with a mean amplitude difference [SD] of 2.51 [1.96] µV on occipito-parietal electrodes between 150 and 200 ms. Following this, the occipito-parietal P2 amplitude was smaller for the Go-uncertain trials relative to the Go-certain trials (mean difference [SD] = 1.15 [1.31] µV between 200 and 300 ms; Figure 4D). These effects collectively constituted a sustained, significant, negative cluster that started in central regions and extended to parieto-occipital electrodes between 70 and 268 ms (sum(t)= -14302, p = 0.0001; cluster 3; see Supplementary Table 2 and Supplementary Figure 5). Additionally, a significant positive cluster between 112 and 242 ms reflected the fronto-temporal counterparts of these effects (sum(t) = 3525, p = 0.006, cluster 2, Supplementary Table 2 and Supplementary Figure 5).

Finally, the central N2 showed a reduced amplitude, with a mean difference [SD] of 1.72 [1.34] µV between 340 and 370 ms, and the occipito-parietal P3 showed an increased amplitude, with a mean difference [SD] of 1.19 [1.52] µV between 350 and 450 ms, for Go-uncertain relative to Go-certain imperative stimuli (Figure 4D). These differences were reflected by a significant, extended, positive cluster spanning over central regions and extending to parieto-occipital electrodes between 260 and 430 ms (sum(t) = 11431, p= 0.0001, cluster 1, Supplementary Table 2 and Supplementary Figure 5).

#### 3.2.2. Effect of reactive inhibition: comparison between Go-uncertain and NoGo trials

As expected, our analysis revealed no significant difference in the ERPs following the cue stimulus for the NoGo and the Go-uncertain trials, where the cue consisted in a red cross in both conditions (Figure 4A).

In contrast, there were differences in ERPs following the imperative stimuli, reflected by several large, significant, positive and negative, clusters (all corrected ps < .05). These effects are detailed below, following the same approach as previously outlined. First, the early central and focal P1 showed reduced amplitude for the NoGo compared to the Go-uncertain trials, with a mean difference [SD] of 1.67µV [2.06] µV between 100 and 130 ms on central electrodes. Concomitantly, the amplitude of the occipital P1 was larger for the NoGo trials than the Go-uncertain trials (mean difference [SD] = 1.52 µV [1.83] µV between 90 and 110ms). Second, the amplitude of the occipito-parietal N1 was larger for the NoGo trials compared to the Go-uncertain stimuli, with a mean difference [SD] of 1.50 [1.23] µV between 150 and 200ms. Third, the amplitude of the N2 component was larger for the NoGo than the Go-uncertain trials, with a mean difference [SD] of 2.28 [2.62] µV on central electrodes between 200 and 300ms. These effects were encompassed collectively in two extended and significant clusters, one negative (sum(t) = -13127, p = 0.0001) and the other positive (sum(t) = 5088, p=0.0001), encompassing the various ERP components (respectively cluster 4 and cluster 1 in Supplementary Table 3; Supplementary Figure 6). Furthermore, a significant positive cluster between 142 and 220 ms delineated the fronto-temporal counterparts of these effects (sum(t) = 3399, p = 0.002, cluster 2 in Supplementary Table 3).

Lastly, we observed two distinct P3 components contingent upon the experimental condition. The P3 was maximal over occipito-parietal electrodes for Go-uncertain trials relative to NoGo trials (mean amplitude differences [SD] = 1.49 [2.35] μV between 350 and 450 ms), whereas it reached its maximum over frontal areas for NoGo trials (mean difference [SD] =2.26 [2.65] μV between 350 and 450 ms). These differences were reflected by two distinct clusters, one negative and one positive, between 300 and 500 ms (sum(t) = - 6916, p = 0.002 and sum(t) = 2968, p = 0.004 respectively; negative cluster 5 and positive cluster 3 in Supplementary Table 3; Supplementary Figure 6).

### 3.3. Time-frequency analysis

#### 3.3.1. Effects of proactive inhibition: comparison between Go-certain and Go-uncertain trials

First, upon computing the TF transform within the time window centered on the cue stimulus, we observed prominent pre-stimulus occipital alpha activity (∼10 Hz) (Figure 5). Using the cluster-based analysis in the pre-stimulus period (between -500 to 0 ms), we found increased alpha band power for the Go-uncertain trials compared to the Go-certain trials (mean difference [SD] = 0.73 [1.66] μV²). This effect manifested in a significant right-lateralized parieto-occipital cluster within the [7-12 Hz] frequency band and [-500 ms; -300 ms] time window (sum(t) = 334, p = 0.004; cluster 1 in Supplementary Table 4).

**Figure 5.**
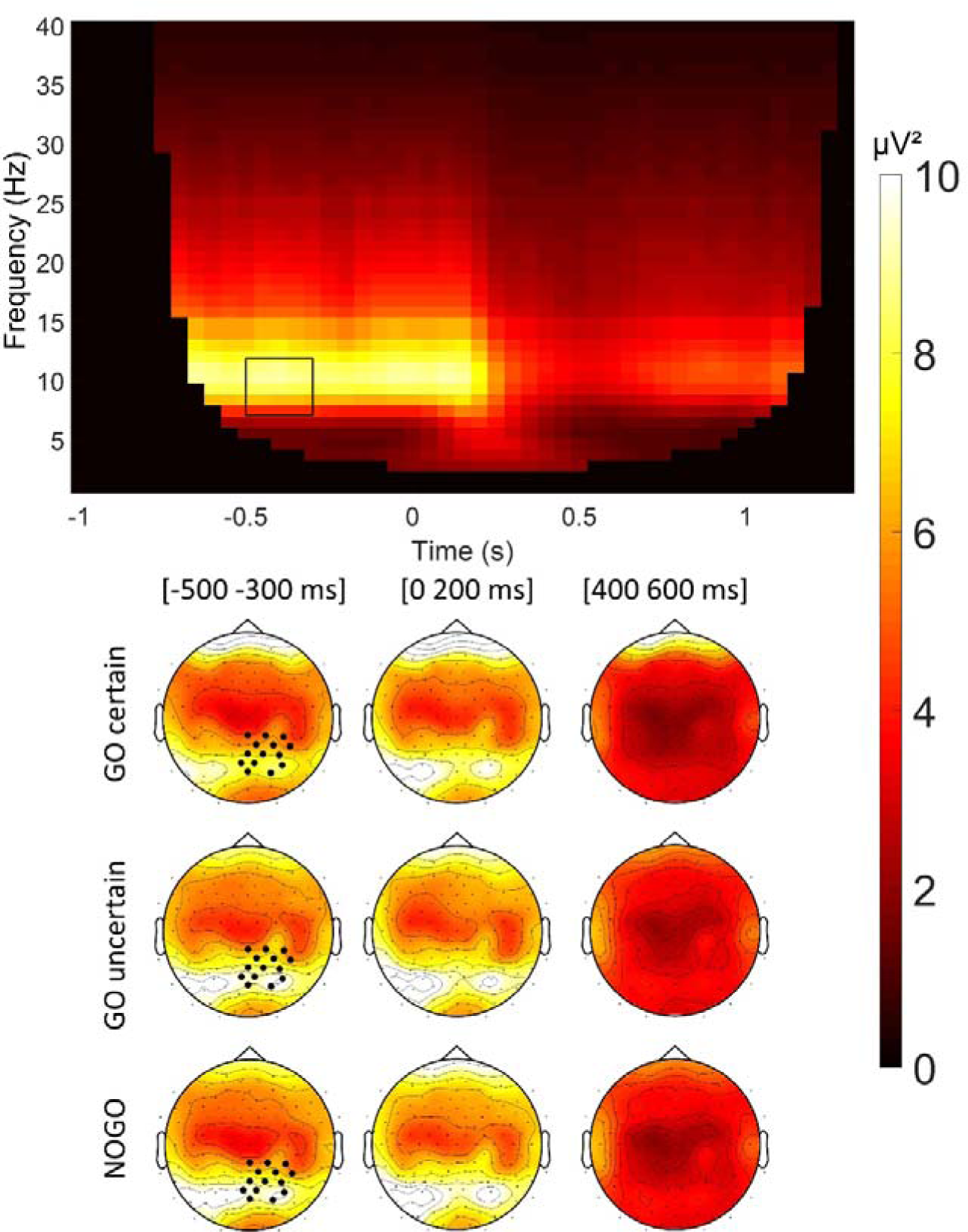
Time-frequency representation of activities in the time period centered on the preparatory stimulus, without baseline correction. <u>Upper part:</u> Grand mean across subjects of the time-frequency representation of the activities before and after the preparatory stimulus onset, without baseline correction. The activities in the TF domain were averaged on the electrodes of the cluster showing a significant difference between the Go-certain and uncertain conditions between -500 and - 300 ms (marked by a black rectangle). These cluster’s electrodes are marked by a black filled circle (•) in the topographical maps underneath the TF representation. <u>Lower part:</u> Topographical maps of activities in the alpha band for the Go-certain, Go-uncertain, and NoGo conditions, averaged between 7 and 12 Hz and in three time windows: [-500; -300ms], [0; 200 ms], [400-600 ms]. These maps represent the grand mean across subjects of the TF data.

Subsequently, we applied baseline correction to the TF data and tested for differences in the post-cue time period. The cluster-based analysis did not reveal any significant difference between Go-uncertain and Go-certain trials in the TF data following the preparatory cue stimulus.

In contrast, significant differences between Go-certain and uncertain trials were observed in the TF domain following the imperative stimulus. We observed increased theta band activity, peaking at 5 Hz, post-imperative stimuli; it was significantly higher for Go-uncertain than Go-certain trials, with a mean difference [SD] of 0.73 [0.81] dB between 0 and 250 ms on occipital electrodes and a mean difference [SD] of 0.86 [1.27] dB between 250 and 300 ms on fronto-central electrodes. This difference manifested in a statistically significant positive cluster starting on parieto-occipital electrodes immediately after the onset of the imperative stimulus. This cluster extended to central and frontal electrodes until 400 ms and it encompassed the alpha band around 100 ms (sum(t) = 2901, p = 0.0001, cluster 2 in Supplementary Table 5).

This effect was followed by a decreased alpha/low beta activity, indicative of desynchronisation within the 8-22 Hz range, which was significantly more pronounced for Go-uncertain trials than for Go-certain trials (mean difference [SD] = 0.53 [0.61] dB). This difference was reflected by a significant negative cluster centered on parieto-central regions between 250 and 500 ms (sum(t) = -460, p = 0.0001, cluster 1 in Supplementary Table 5) (Figure 6).

**Figure 6.**
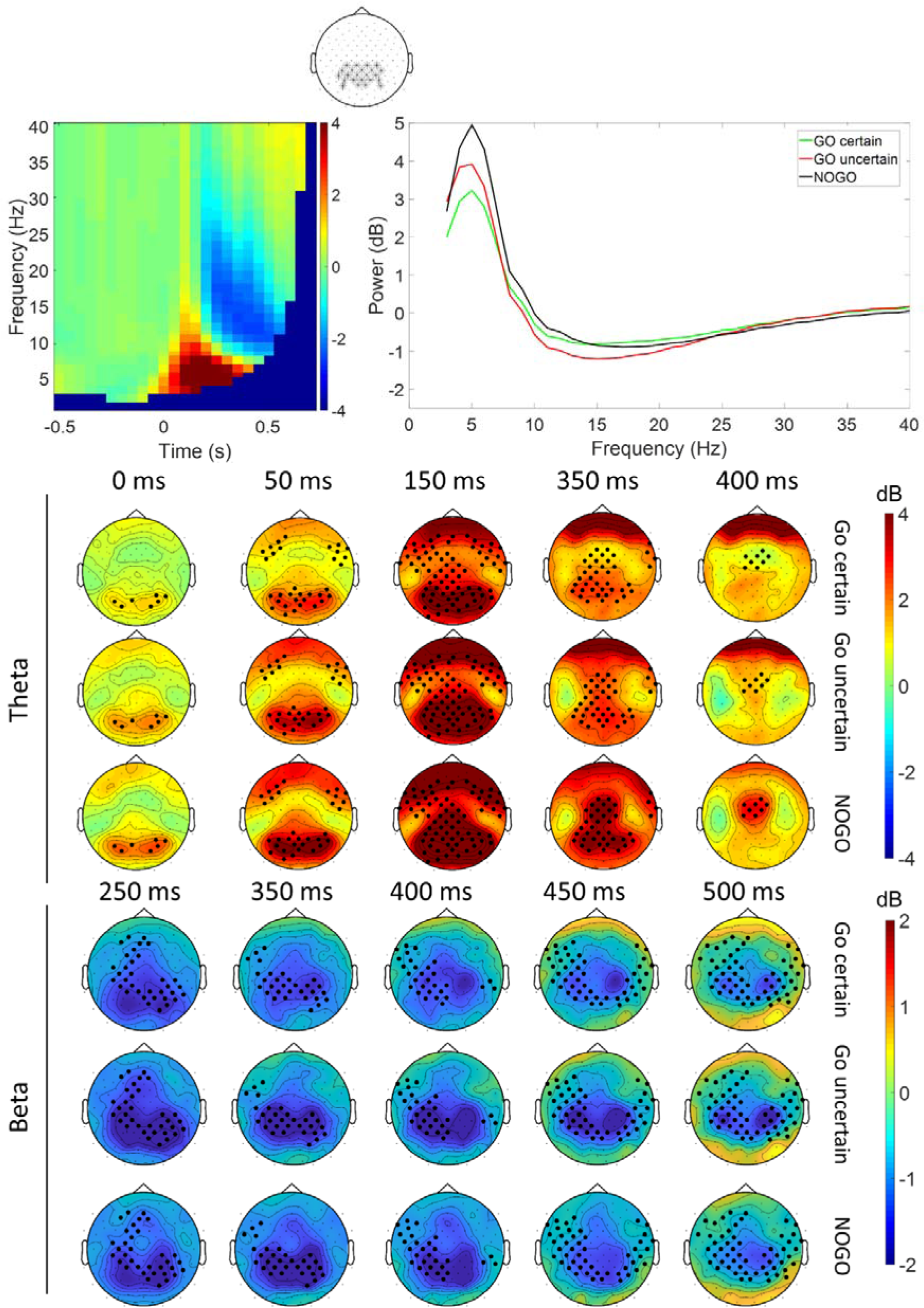
Time frequency representation of activities in response to the imperative stimulus. <u>Upper part:</u> On the left, grand mean across subjects of the time-frequency representation of the activities (in dB) in response to the imperative stimulus, here illustrated in the Go-uncertain condition. On the right, power spectrum (computed between 0 and 500 ms) of the activities between 3 and 40 Hz in the Go-certain (in green), the Go-uncertain (in red) and the NoGo (in black) conditions. Both figures were obtained by averaging TF activities from the electrodes highlighted by an asterisk (*) in the upper map inset. <u>Lower part:</u> Grand mean across subjects of the topographical maps of TF activities (in dB) in the theta (3-7 Hz) and low beta (13-21 Hz) bands, at 0, 50, 150, 350, and 400 ms, for the Go-certain, Go-uncertain, and NoGo conditions. On each map, the cluster’s electrodes from the comparison between NoGo and Go-uncertain conditions, in the considered frequency band and at the illustrated time point, are highlighted with a ‘•’.

#### 3.3.2. Effects of reactive inhibition: comparison between Go-uncertain and NoGo trials

The cluster-based analysis of NoGo versus Go-uncertain conditions in the TF domain revealed significant differences encompassing all frequency bands of interest within a single large statistically significant cluster in the time period following the imperative stimulus (sum(t) = 11022, p = 0.0001; cluster 1 in supplementary Table 6). More specifically, the theta synchronisation that started immediately after the imperative stimulus onset on occipital electrodes and extended to parietal and frontocentral electrodes until 400 ms, was more pronounced for the NoGo than the Go-uncertain trials, with a mean difference [SD] of 0.76 [0.70] dB on occipito-parietal electrodes between 0 and 250 ms and of 1.36 [1.26] dB on fronto-central electrodes between 250 and 300 ms (Figure 6). This effect extended into the alpha band over occipital and central electrodes between 50 and 200 ms (mean difference [SD] = 0.78 [0.80] dB). Then, desynchronisation started in the alpha band (8-12 Hz) on bilateral parietal electrodes and extended to fronto-central electrodes until 450 ms (mean difference [SD] = 0.60 [0.81] dB between 250 and 450 ms); it extended to the low beta band (13-21 Hz) over the centro-parietal electrodes between 100 and 300 ms (mean difference [SD] = 0.34 [0.57] dB). This alpha/low beta desynchronisation was larger for the Go-uncertain than NoGo trials (Figure 6).

## 4. DISCUSSION

This study investigated the impact of proactive and reactive inhibition on gait initiation, using a modified Go/NoGo task. We demonstrated the presence of proactive inhibition during gait initiation in the context of uncertainty in healthy adults, with heightened reaction times in Go-uncertain trials compared to Go-certain trials. At the electrophysiological level, differences between Go-uncertain and Go-certain trials manifested in alpha synchronisation before the preparatory cue stimulus onset and in a larger centro-parietal P3 response to the preparatory cue. Notably, uncertainty had a marked influence on the ERPs and TF activities following the imperative stimulus. Go-uncertain stimuli elicited smaller central P1, larger occipito-parietal N1, smaller occipito-parietal P2 and central N2, larger occipito-parietal P3, increased theta band synchronisation and alpha/low beta band desynchronisation, compared to Go-certain imperative stimuli. These activities were further modulated by reactive inhibition, with reduced central P1, larger occipito-parietal N1 and central N2, smaller occipito-parietal P3, increased theta synchronisation, and reduced alpha/low beta desynchronisation in response to NoGo relative to Go-uncertain imperative stimuli. Additionally, the NoGo stimuli elicited a distinct frontal P3 response.

### 4.1. Behavioural signature of proactive inhibition

At the behavioural level, as expected, we found an increased APA RT, and consequently an increased FO reaction time, in the Go-uncertain relative to the Go-certain conditions. This is in line with the results previously obtained by Albares et al. (2014) and extend these results to gait initiation. On the other hand, the APA duration and the other gait initiation parameters were not significantly modulated by the Go/NoGo uncertainty context. Thus, the uncertainty of the context of gait initiation selectively affected movement preparation, without any significant effect on movement execution. This contrasts with some previous studies that showed an influence of executive processes such as attention on the timing of postural adjustments in different tasks (Sparto et al., 2013; Tard et al., 2013). It suggests that the influence of context on movement execution may be dependent on the executive processes involved in the task.

### 4.2. Proactive inhibition in the time period centered on the preparatory stimulus: involvement of attention-related processes?

At the electrophysiological level, uncertainty was associated first with heightened occipito-parietal alpha activity prior to the preparatory stimulus onset. Synchronisation of brain activities in the alpha band has been associated with inhibition in the region concerned, particularly in relation to attention suppression mechanisms (Haegens et al., 2011; Morrow et al., 2023; Sauseng et al., 2009). In our study, for Go-uncertain trials, the subject had to wait for the imperative stimulus to determine which motor program to initiate; hence the preparatory cue stimulus was less informative than for the Go-certain trials, where the preparatory stimulus indicated upcoming gait initiation. Thus, it may be suggested that, in the condition of uncertainty, the subjects implemented inhibition of the processing of the preparatory stimulus. The fact that this alpha synchronisation was observed prior to the preparatory stimulus onset indicates that it was a block-related cognitive strategy, which was allowed by the presentation of Go-certain and Go-uncertain trials in separate blocks. In addition, we observed a larger occipito-parietal P3 in response to the cue stimuli for Go-uncertain relative to Go-certain conditions. This could reflect the need for increased attention in the uncertain context, in order to prepare for processing and discriminating the incoming imperative stimulus (for review see Verleger, 2020; Polich, 2007; Tard et al., 2013), and thus reflecting “proactive inhibition” where subjects had to wait for the imperative stimulus for motor decision and planning. In this respect, proactive inhibition processes may at least partly overlap with attention-related processes (Perri, 2020; see also Tatz et al., 2021).

### 4.3. Proactive inhibition and reactive inhibition following the imperative stimulus: same or different inhibitory processes?

We found extended and overlapping modulations of the brain activities in response to the imperative stimuli for Go-uncertain versus Go-certain trials (proactive inhibition) and for NoGo versus Go-uncertain trials (reactive inhibition), in both ERP and TF analyses.

First, we observed an early focal P1 that peaked around 100 ms on midline central electrodes. To the best of our knowledge, this focal, central P1 has not been described before. It was greater for the Go-certain than the Go-uncertain stimuli and virtually absent for the NoGo stimuli. It could therefore be linked to the early phase of gait initiation, that is, to the APA phase. Interestingly, an early negative component peaking over central regions around 100 ms has previously been reported in response to exogenous postural imbalance (Marlin et al., 2014; Mierau et al., 2015; Mochizuki et al., 2017; Varghese et al., 2014). In our study, the postural imbalance was endogenous, voluntary, and planned, since it was related to the APAs for gait initiation. Thus, it is possible that both types of early activities were related to postural control, with the negativity reflecting inhibition of or counter-activation to exogenously imposed postural imbalance and the positivity reflecting endogenous activation of postural control associated with the APAs. The decreased amplitude of the central P1 for the Go-uncertain trials may reflect incomplete lifting of proactive inhibition in this early time window for this condition, which was associated with delayed APAs onset. In line with this hypothesis, the early postural negativity has been reported to be less prominent when the exogenous postural imbalance can be anticipated, in comparison to the case of unpredictable perturbation (Mochizuki et al., 2017).

Following the P1, we observed a large occipito-parietal N1, which peaked around 170 ms and was larger for the Go-uncertain versus Go-certain trials and for the NoGo versus Go trials. This is partly reminiscent of the dMF170 of Albares et al (2014) obtained by contrasting Go-certain and uncertain conditions (see also De Blasio & Barry, 2013) and whose amplitude on FCz was negatively correlated with the upper limb reaction times to Go stimuli. In the present study, it is likely that the effects on occipito-parietal N1 that we observed using a classical ERP analysis encompassed modulations of the so-called dMF170. Accordingly, the cluster-based statistical analysis showed effects (aka. clusters) extending to fronto-central electrodes, with polarity reversal on anterior fronto-temporal regions (see Supplementary Figure 5). That said, in our study, the occipito-parietal N1 was modulated in both Go-uncertain versus Go-certain and NoGo versus Go-uncertain trial comparisons, whereas it was only present for the comparison between Go trials in the study of Albares et al. This apparent discrepancy may be explained by several experimental differences between the two studies. First, we used a blocked design, where certain and uncertain conditions were presented in separate blocks, whereas these conditions were intermixed in Albares et al. (2014), thus favoring different cognitive strategies (Coxon et al., 2007). Second, our task always started with a block of Go-certain trials, then followed by 50% / 50% of mixed Go-uncertain and NoGo trials. Although this 50/50 probability of Go/NoGo trials may be considered as not optimal for the elicitation of selective reactive inhibition in response to the NoGo stimuli, our design starting with a Go-certain block and including interleaved Go-certain and -uncertain blocks put some emphasis on selective reactive inhibition in response to NoGo stimuli. Lastly, and importantly, gait initiation is a much more complex motor task than a simple button press, thus requiring selection between two alternative responses, namely initiating gait or maintaining an immobile standing position. It is therefore possible that our protocol allowed showing evidence for the early inhibitory processes related not only to proactive inhibition and/or nonselective reactive inhibition but also to selective reactive inhibition. Altogether, our results support the view that the activities in the early time window of the N1 were associated with both proactive and reactive inhibition processes (De Blasio & Barry, 2013), reflecting early automatic inhibitory processes associated with the overall level of inhibition involved in the task (Albares et al., 2014; Barry et al., 2016).

In the TF domain, theta-band activity in response to the imperative stimuli showed a similar pattern of modulation with increasing theta power for Go-certain, then Go-uncertain, and finally NoGo conditions. This is in line with the proposed role of theta band activities in response control and conflict monitoring (Cohen & Donner, 2013; De Blasio & Barry, 2013; J. Kaiser et al., 2022). Interestingly, this theta activity was maximal over posterior, occipito-parietal regions around 150 ms and extended then to fronto-central regions until about 400 ms. Theta band activity has been related to communication between brain regions (Anderson et al., 2010; Cavanagh & Frank, 2014; Karakaş, 2020; Lisman & Jensen, 2013; Zhang et al., 2018). This suggests that the theta band-activity could reflect communication between sensory areas processing visual information and motor areas involved in our gait initiation task.

We also observed a P2 that followed the N1 on occipito-parietal regions and was greater in response to Go-certain than Go-uncertain stimulus. The functional significance of this component is unclear. It may reflect some advanced perceptual processing of stimuli, related for instance to stimulus identification (Lindholm & Koriath, 1985; Omoto et al., 2010) and affected by repetition, probability, and attention (Phillips & Takeda, 2009; Qian et al., 2012; Tremblay et al., 2001). It could reflect inhibition of sensory information processing (Hegerl & Juckel, 1993). The effect of uncertainty on P2 might therefore be related to the suppression of extensive processing of the imperative stimulus in the Go-certain condition, where only the detection of the stimulus was necessary, allowing prioritization of the motor response.

### 4.4. The N2/P3 complex and alpha/low beta activities are associated with inhibition and motor control

The late component of the ERPs, i.e. the N2/P3 complex, showed differences among the three conditions. The N2/P3 components have long been associated with inhibitory processes, particularly reactive inhibition (e.g., Maguire et al., 2009; Nguyen et al., 2016; Oddy & Barry, 2009), but also with various other processes, such as attention, memory, cognitive control, etc. in the context of Go/NoGo paradigms, rendering their interpretation quite complex (Asanowicz et al., 2020; Falkenstein et al., 1999; Folstein et al., 2008; Friedman et al., 2001; Huster et al., 2013; Judah et al., 2013; Polich, 2007; Verleger, 2020). The late latency of these activities makes them unlikely to reflect purely inhibitory processes (Albares et al., 2015). The fact that the N2 wave was less prominent in the Go-uncertain trials, but with similar amplitude in the Go-certain and NoGo trials, suggests that it might be associated with the continuation of a planned motor program. In the Go-certain trials, the subject had to move in each trial and only waited for the signal to initiate this pre-activated motor plan. In the NoGo trials, the subject had to continue the standing-still motor plan. In contrast, in the Go-uncertain trials, the subject had to change the motor plan from the standing immobile position to gait initiation in response to the imperative stimulus. The reduced N2 may reflect this motor plan switch.

The N2 wave was followed by occipito-parietal P3 in response to Go stimuli and frontal P3 in response to NoGo stimuli. The occipito-parietal P3 was more prominent in response to Go-uncertain than Go-certain stimuli. In continuity with the preceding N2, it may reflect the sustained action control required for initiating gait, with increased need for online action monitoring in the Go-uncertain relative to Go-certain conditions (Beste et al., 2012; Walentowska et al., 2016). The P3 component in response to NoGo stimuli had a distinct, frontal topography, typical of the so-called NoGo P3, which has been extensively studied in relation to reactive inhibition. This NoGo P3 has long been associated with selective reactive inhibitory processes triggered by the NoGo stimuli (Bokura et al., 2001; Falkenstein et al., 1999; Hervault et al., 2022; Smith et al., 2008, 2010). However, its late latency rather suggests that it is associated with post-inhibition processes (Albares et al., 2015; Huster et al., 2020). The present data are consistent with this view in that Go-uncertain occipito-parietal and NoGo frontal P3 may reflect the distinct action control required for gait initiation and standing still posture maintenance, respectively.

The N2/P3 complex was concomitant with alpha/low beta desynchronisation in the TF domain. This desynchronisation peaked on bilateral parietal and central regions and was more pronounced in response to the Go-uncertain relative to the Go-certain and NoGo imperative stimuli. P3 amplitude and latency have been associated with alpha desynchronisation (Polich, 2011, 2007). Moreover, alpha/beta desynchronisation in motor regions has been associated with movement preparation and execution (Jasper & Andrews, 1936; Kilavik et al., 2013; Pfurtscheller & Lopes da Silva, 1999). It may reflect the increased motor preparation and initiation required for gait initiation in Go-uncertain trials, in order to counteract the ongoing inhibitory process associated with uncertainty (Gwin & Ferris, 2012; Kilavik et al., 2013; Zaepffel et al., 2013).

### 4.5. Limitations

The present study has several limitations. First, it is based on a relatively small sample of subjects, which excludes a strong generalisation of our results. Statistical power analysis indicated that with 23 subjects, effects with standardized size (Cohen’s dz) of 0.61 could be detected (with a power of 0.8 and a p-value of .05; sensitivity analysis for two-tailed paired t-tests in G*Power 3.1.9.2), corresponding to medium size effects. Importantly, we obtained the expected behavioural effect of Go-uncertainty with increased RT for all but one individual subject, with also very few omission and commission errors. It is also noteworthy that there were some partial errors in the form of lateral shifts of the CoP without ensuing foot-off in response to the NoGo stimuli. This supports the view that our experimental protocol elicited robust effects related to inhibitory processes, including both proactive and reactive processes during gait initiation, even if the 50/50 probability of Go/NoGo trials during Go-uncertain blocks is atypical and can be considered as not optimal to elicit reactive inhibition. Although the studies of reactive inhibition typically include more frequent Go than NoGo stimuli (e.g., Benikos et al., 2013; Nguyen et al., 2016), this is not a necessary condition (Albares et al., 2014; Barry et al., 2014; De Blasio & Barry, 2013; Gajewski & Falkenstein, 2013; Karamacoska et al., 2018; Liebrand et al., 2017; Schmiedt-Fehr et al., 2016), as confirmed by the present study. The inclusion of Go-certain blocks also favoured gait initiation as the prepotent response in our protocol.

Go-certain and -uncertain conditions were well comparable, because both entailed the same behavioural response (aka. gait initiation). In contrast, there was an intricacy of both inhibitory and motor execution processes in the comparison between Go-uncertain and NoGo trials. We interpreted our results carefully in keeping with this limitation, considering both inhibitory and motor accounts of our findings (Figure 7). Including the Go-certain condition, with minimal inhibitory activity, allowed us to partly disentangle both accounts.

**Figure 7.**
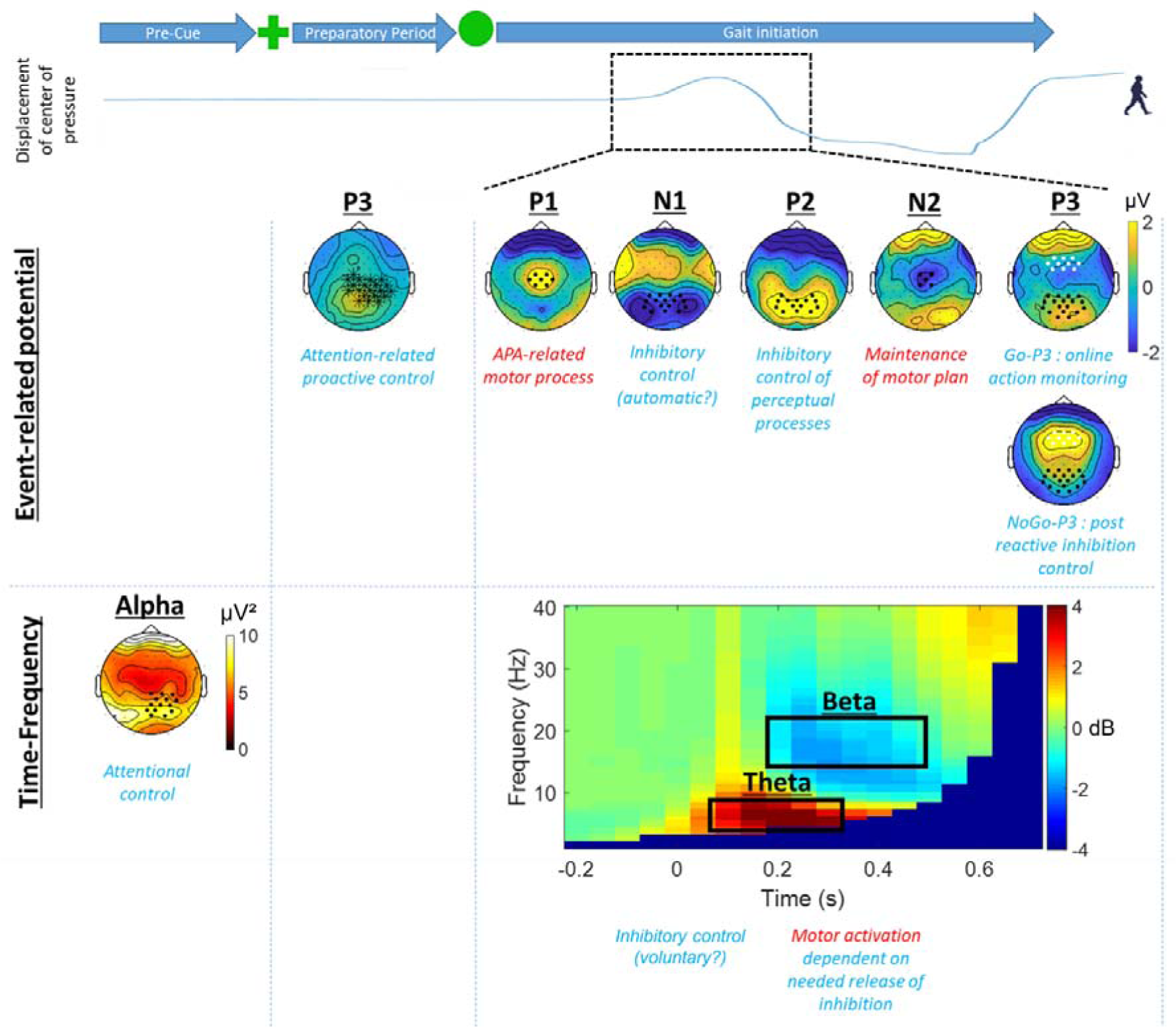
Synthetic overview of the findings and their interpretation. In our Go/NoGo gait initiation task including a Go certain condition, we found extended effects on multiple EEG activities in the ERP and TF domains. We summarise here the main interpretations proposed for these effects in term of inhibitory and more generally executive processes (in turquoise blue) and in terms of motor processes (in red).

The EEG data associated with gait contained large artefacts and one may wonder if this may have affected our results. In this regard, we implemented thorough reviewing and correction of data to avoid any confounding effect of artefacts (see Methods). Moreover, we limited our time window of statistical analysis to 500 ms after the imperative stimulus. It may also be noted that we used a 1-Hz high-pass filter for data pre-processing. While this may attenuate late latency components such as the P3, it has a short time constant of ∼160 ms, avoiding propagation of distortion due to any remaining transient artefact related to gait into the data time window of interest. This choice was made following Gwin et al. (2010), but it may limit comparison with the previous studies that used high-pass filters with lower cut-off (e.g., Delval et al., 2016; Wagner et al., 2016).

In this study, we adopted classical ERP and TF analysis approaches. This allowed us to show the sustained influence of inhibitory processes on electrophysiological activities associated with gait initiation. It is likely that the effects we observed involved multiple underlying sources in candidate regions of the frontal and parietal lobes known to be involved in motor and inhibitory control (Banich & Depue, 2015; Cavanagh & Frank, 2013; Diesburg & Wessel, 2021; Osada et al., 2019; Schall et al., 2017; Wiecki & Frank, 2013). Source separation and localization techniques may be used in the future to further characterize these brain sources. One may also wonder if there was any brain-behaviour correlation in our data. We performed such correlation analysis on an exploratory basis, but they did not yield any significant result, probably due to our relatively small sample size. Lastly, additional psychometrics/cognitive tests might have allowed to further link our data with executive functions for better understanding the inhibitory processes in the context of walking.

### 4.6. Conclusion

Proactive and reactive inhibitory processes appear as two tightly interlinked cognitive processes in gait initiation. They are associated with extended and overlapping modulations of electrophysiological activities, which reflect the continuous control of information processing and action during gait initiation. Our results emphasize the interplay between the release of proactive inhibition, the setting of reactive inhibition, action selection, and movement initiation in relation with the uncertainty of gait initiation context. These results may be important for understanding difficulties of gait initiation in pathological context, such as Parkinson’s disease. Future studies will allow us to examine the alterations of the activities uncovered here in this pathology, and to further explore the neural basis of inhibitory processes during gait initiation, for example using source localization and functional connectivity approaches.

## Financial disclosure in the last 12 months

D Ziri, C Olivier, P Boulinguez, H.Gunasekaran, B Lau, L Hugueville, N George have nothing to disclose.

M.L. Welter has received consultancy fees from Medtronic, MindMaze France, and Boston Scientific

## Acknowledgments

We thank all the participants in this study. We also thank the administrative staff of the Clinical Investigation Centre and Marion Albares for her help in the conception of this study. We thank Edward Soundaravelou and Clara Mouillevois for their help in data acquisition. This study was supported by a grant from the Agence Nationale de la Recherche (project LOCOMOTIV, nb. ANR-16-CE37-0007), and ‘Investissements d’avenir’ (ANR-10-IAIHU-06). D.Ziri was funded by France Parkinson (contract number: 2203027NA). The funding sources did not have any involvement in either study design, data collection, analysis, interpretation, or the writing of the report and decision to submit the article for publication.

## Declaration of authors’ contribution

Research conception and design: M-L Welter, P Boulinguez, B Lau; Participants’ recruitment and inclusion: M-L Welter; Data acquisition: D Ziri, C Olivier, L Hugueville, with contribution of N George. Data analysis: D Ziri, C Olivier, L Hugueville, H Gunasekaran, with the supervision of N George, M-L Welter. Result interpretation: D Ziri, N George, M-L Welter; writing of the article: D Ziri, N George, M-L Welter; article revision: all co-authors. All authors approved the final version of the manuscript.

