## Supplementary Material for "Inhibitory control of gait initiation in humans: an electroencephalography study"

**Supplementary Figure 1**: Example CoP trajectories on the medio-lateral axis during a Go certain trial (A) and during two NoGo trials, one without (B) and the other with (C) partial error. (A) For each trial, the medio-lateral position of the CoP (in mm) was plotted along time (in s) and markers were manually placed on the events of interest. Time 0 marked the start of the trial by the experimenter (remind that each trial was started manually once the subject was settled, immobile, at the starting point on the force platform). In the illustrated trial, the preparatory cue stimulus appeared 1s after the trial start and it was followed by the imperative Go stimulus after 1s. One can see that the CoP line was flat before the Go stimulus, indicating that the subject did not move (only some weak natural oscillations can be detected). The first displacement of the CoP corresponded to the beginning of the APA (red vertical line); it was followed by the first foot off (vertical blue line) and the completion of the first step (magenta vertical line). Then, the second foot off took place (green vertical line) followed by the second step (vertical turquoise blue line). The heel off (vertical black line) preceding the first foot off was marked using the visualisation of heel markers on the Vicon Nexus 2.10.3 system. (B) Illustration of the medio-lateral position of the CoP (in mm) along time (in s) during a NoGo trials where no partial error was detected. One can see that the CoP line remained flat across the trial apart from weak natural oscillations seen on every trial. (C) Illustration of the medio-lateral position of the CoP (in mm) along time (in s) during a NoGo trial with a partial error. This partial error can be seen in the form of a well detectable oscillation of the CoP in response to the NoGo stimulus not followed by foot-off.

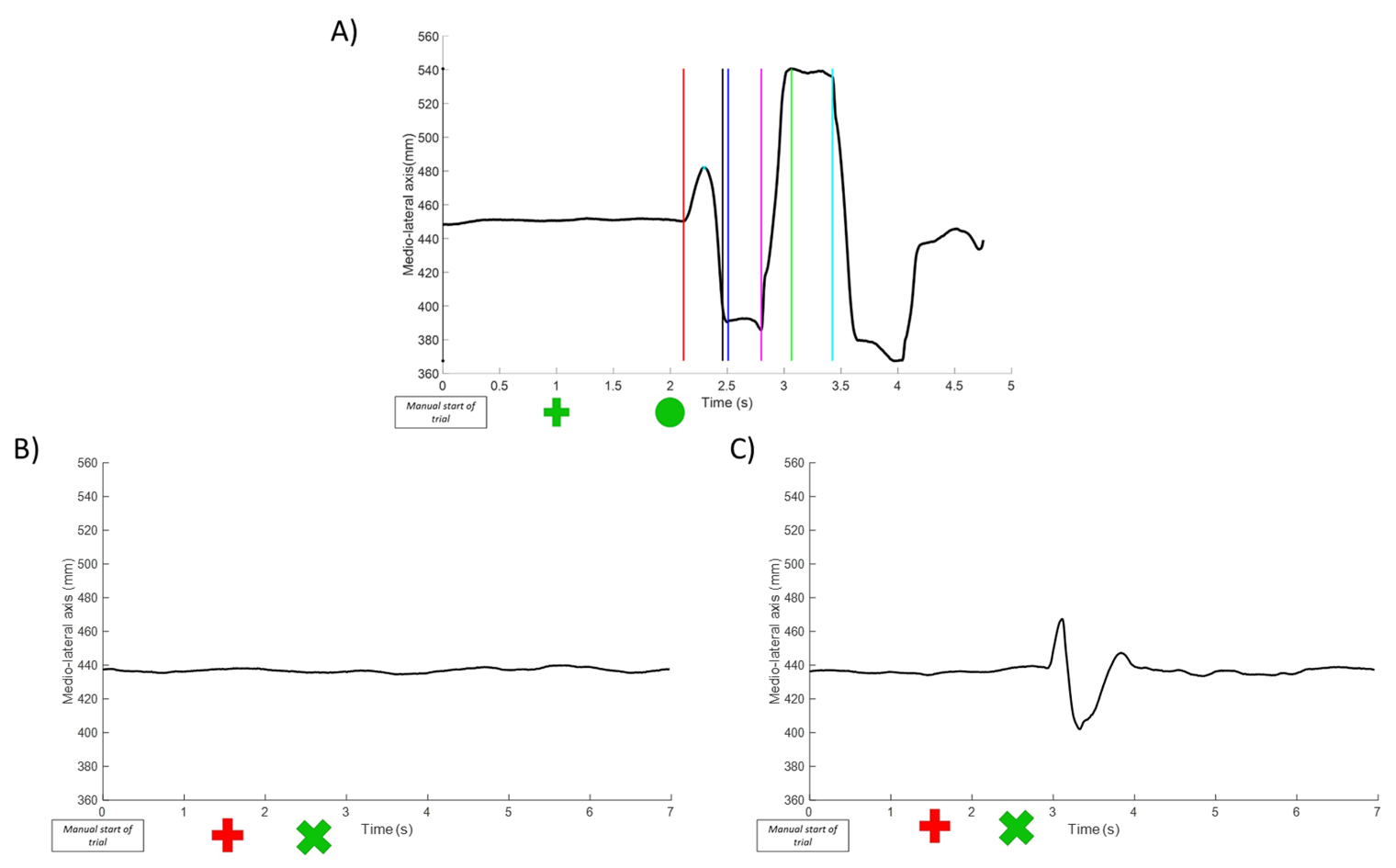

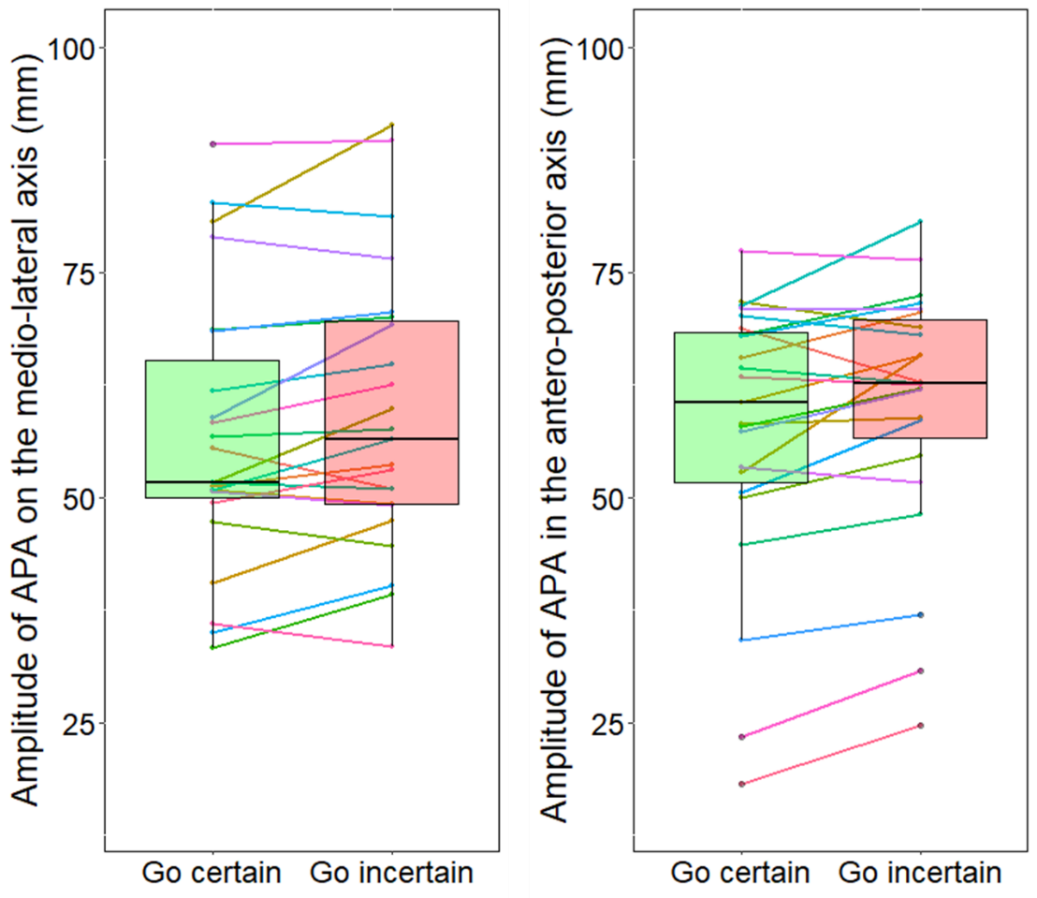
**Supplementary Figure 2**: Amplitude of APAs in the Go certain (in green) and the Go uncertain (in red) conditions

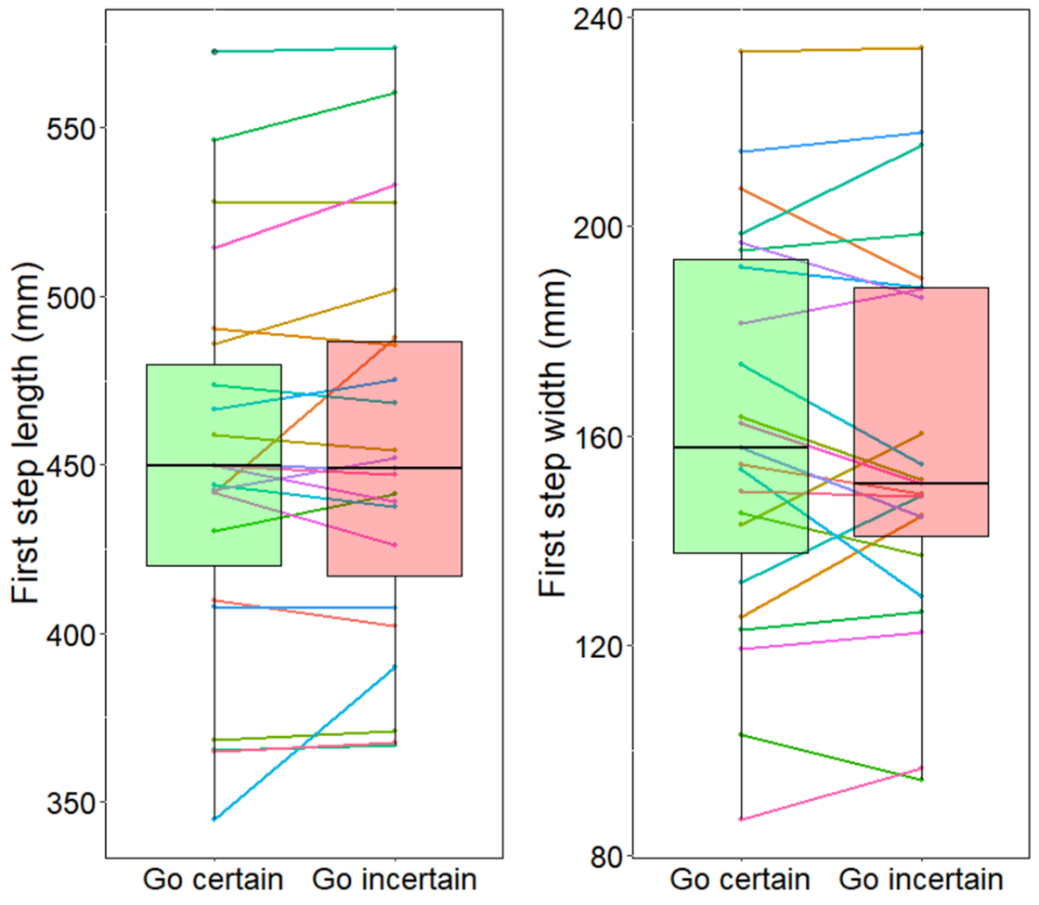
**Supplementary Figure 3:** First step parameters in the Go certain (in green) and the Go uncertain (in red) conditions

**Supplementary Table 1:** ERP cluster obtained by comparison between the Go certain and Go uncertain conditions in response to the preparatory stimulus

| Cluster n° | Electrodes | Time (ms) | Cluster statistic [sum(t)] | Corrected p value |
| --- | --- | --- | --- | --- |
| 1 | FCC1h, C1, Cz, C2, C4, CCP1h, CCP2h, CCP4h, CCP6h, CP1, CPz, CP2,  CP4, CP6, CPP2h, CPP4h, CPP6h, Pz, P2, P4 | 480 - 510 | 584 | 0.037 |

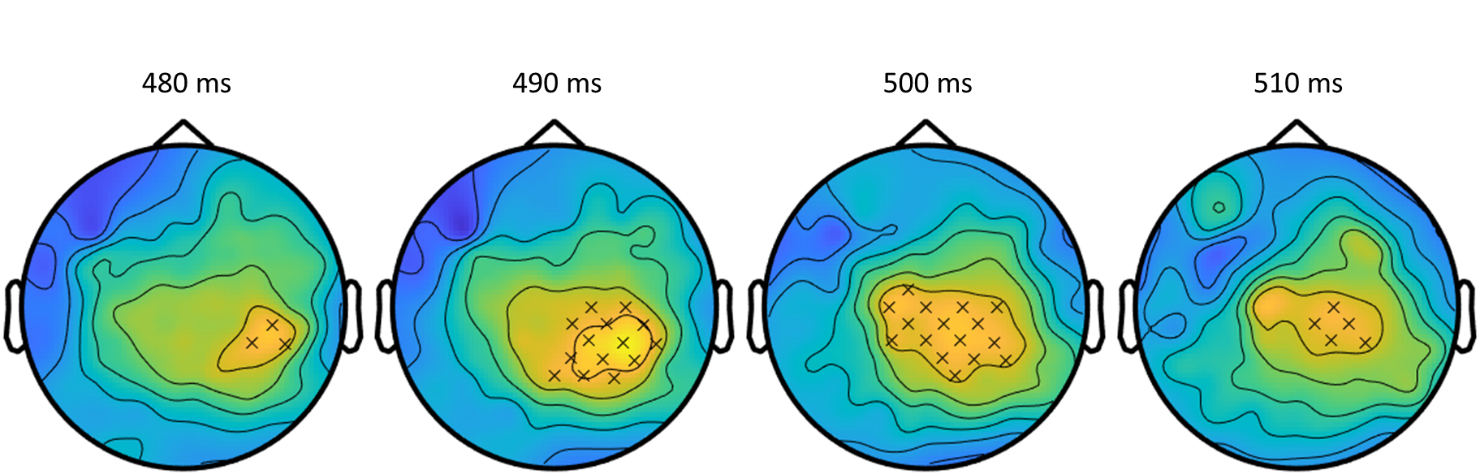
**Supplementary Figure 4:** Illustration of the dynamics across time of the cluster in Supplementary Table 1

**Supplementary Table 2:** ERP clusters obtained by comparison between the Go certain and Go uncertain conditions in response to the imperative stimulus

| Cluster n° | Electrodes | Time (ms) | Cluster statistic [sum(t)] | Corrected p value |
| --- | --- | --- | --- | --- |
| 1 | Fz, F2, FFC5h, FFC1h, FFC2h, FFC4h, FFC6h, FC5, FC3, FC1, FCz, FC2, FC4, FC6, FCC5h, FCC3h, FCC1h, FCC2h, FCC4h, FCC6h, C5, C3, C1, Cz, C2, C4, TTP7h, CCP5h, CCP3h, CCP1h, CCP2h, CCP4h, CCP6h, CP5, CP3, CP1, CPz, CP2, CP4, CP6, TPP7h, CPP5h, CPP3h, CPP1h, CPP2h, CPP4h, CPP6h, P5, P3, P1, Pz, P2, P4, P6, PPO5h, PPO1h, PPO2h, PPO6h, PO3, POz, PO4 | 260-430 | 11431 | 0.0001 |
| 2 | AF3, AFz, AF4, AFF5h, AFF1h, AFF2h, AFF6h, F7, F5, F3, F1, Fz, F2, F4, F6, F8, FFT9h, FFT7h, FFC6h, FFT8h, FFT10h, FT7, FC6, FT8, FTT9h, FTT10h, T7, T8 | 112-242 | 3525 | 0.006 |
| 3 | FFC3h, FC3, FC1, FCz, FC4, FCC5h, FCC3h, FCC1h, FCC2h, FCC4h, FCC6h, C5, C3, C1, Cz, C2, C4, CCP5h, CCP3h, CCP1h, CCP2h, CCP4h, CCP6h, TTP8h, CP5, CP3, CP1, CPz, CP2, CP4, CP6, TPP7h, CPP5h, CPP3h, CPP1h, CPP2h, CPP4h, CPP6h, P7, P5, P3, P1, Pz, P2, P4, P6, PPO5h, PPO1h, PPO2h, PPO6h, PO7, PO3, POz, PO4, PO8, POO1, POO2, O1, Oz, O2 | 70-268 | -14302 | 0.0001 |

**Supplementary Figure 5:** Illustration of the dynamics across time of the clusters in Supplementary Table 2

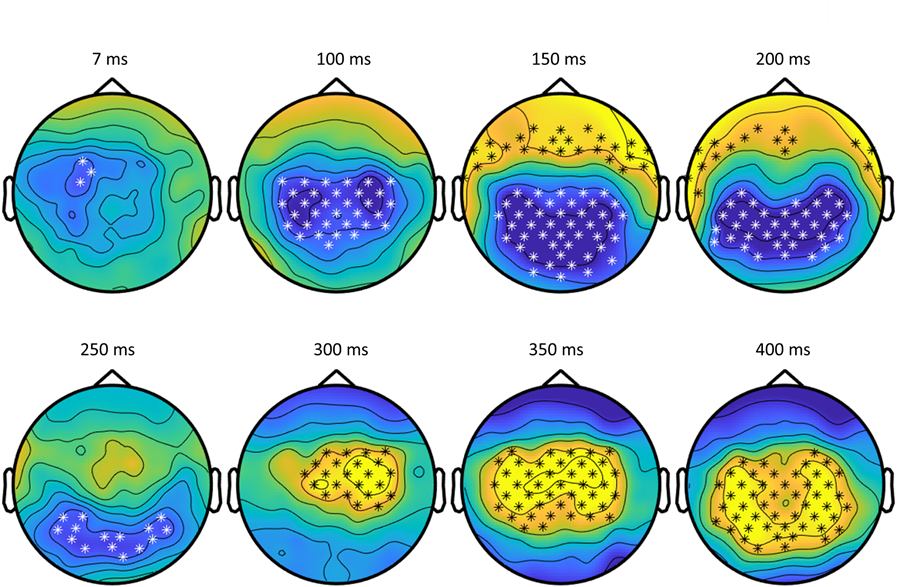

**Supplementary Table 3:** ERP clusters obtained by comparison between the Go uncertain and NoGo conditions in response to the imperative stimulus

| Cluster n° | Electrodes | Time (ms) | Cluster statistic [sum(t)] | Corrected p value |
| --- | --- | --- | --- | --- |
| 1 | CCP5h, CP5, CP3, CP4, CP6, TP8, TPP7h, CPP5h, CPP3h, CPP1h, CPP4h, CPP6h, TPP8h, TPP10h, P7, P5, P3, P1, Pz, P2, P4, P6, P8, PPO5h, PPO1h, PPO2h, PPO6h, PO7, PO3, POz, PO4, PO8, POO9h, POO1, POO2, O1, Oz, O2 | 0-136 | 5088 | 0.0001 |
| 2 | AF3, AFz, AF4, AFF5h, AFF2h, AFF6h, F7, F5, F4, F6, F8, FFT9h, FFT7h, FFC6h, FFT8h, FFT10h, FT7, FC5, FC6, FT8, FTT9h, FTT7h, FCC6h, FTT8h, FTT10h, T7, C5, C6, T8, TTP7h, CCP6h, TTP8h, CP6, TP8 | 142 - 220 | 3399 | 0.002 |
| 3 | AFF1h, AFF2h, F3, F1, Fz, F2, F4, FFC5h, FFC3h, FFC1h, FFC2h, FFC4h, FFC6h, FC3, FC1, FCz, FC2, FC4, FCC3h, FCC1h, FCC2h, FCC4h, C3, Cz, C2, CCP3h | 372-498 | 2968 | 0.004 |
| 4 | AFF1h, AFF2h, F3, F1, Fz, F2, F4, FFT9h, FFT7h, FFC5h, FFC3h, FFC1h, FFC2h, FFC4h FFC6h, FFT8h, FT7, FC5, FC3, FC1, FCz, FC2, FC4, FC6, FTT9h, FCC5h, FCC3h, FCC1h, FCC2h, FCC4h, CC6h, C3, C1, Cz, C2, C4, CCP5h, CCP3h, CCP1h, CCP2h, CCP4h, CP1, CPz, CP2, CPP5h, CPP3h, CPP1h, CPP2h, CPP4h, TPP8h, P5, P3, P1, Pz, P2, P4, P6, P8, PPO5h, PPO1h, PPO2h, PPO6h, PO7, PO3, POz, PO4, PO8, POO9h, POO1, POO2, O1, O2 | 12-304 | -13127 | 0.0001 |
| 5 | C3, C6, TTP7h, CCP5h, CCP3h, CCP6h, TTP8h, TP7, CP5, CP3, CP1, CP4, CP6, TP8, TPP7h, CPP5h, CPP3h, CPP4h, CPP6h, TPP8h, TPP10h, P7, P5, P3, P1, Pz, P2, P4, P6, P8, PPO5h, PPO1h, PPO2h, PPO6h, PO7, PO3, POz, PO4, PO8, POO9h, POO1, POO2, O1, Oz, O2 | 289-446 | -6915 | 0.002 |

**Supplementary Figure 6**: Illustration of the dynamics across time of the clusters in Supplementary Table 3

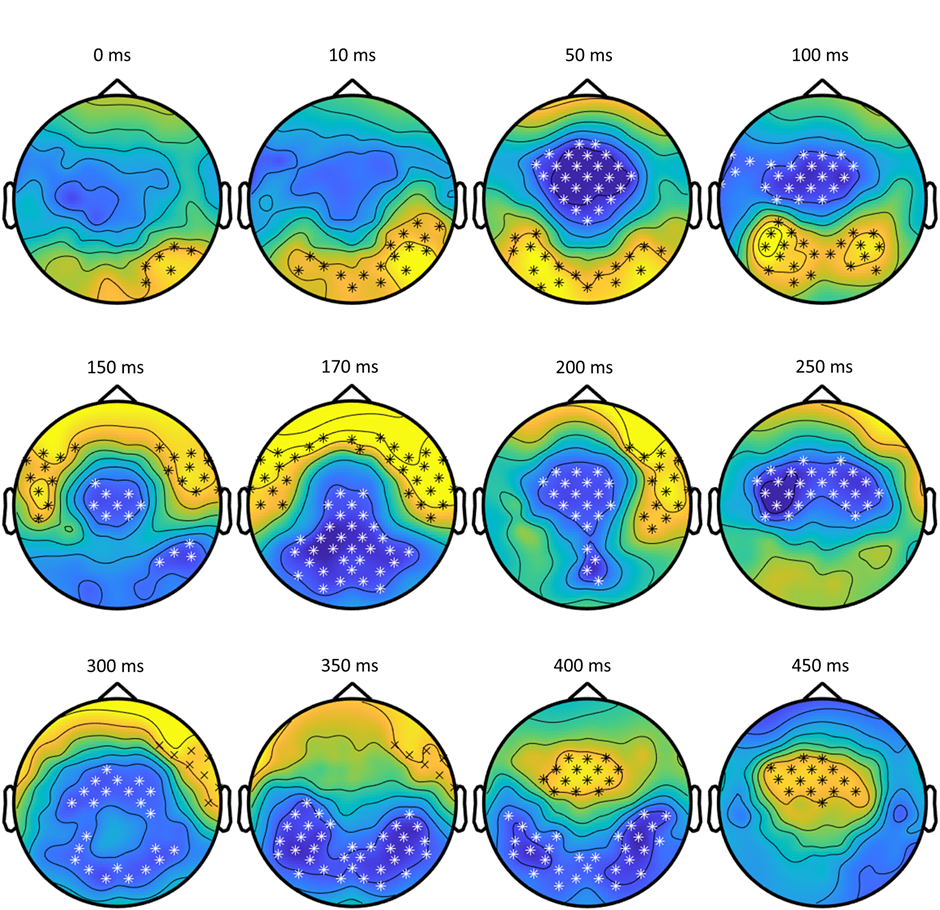

**Supplementary Table 4:** TFR cluster obtained by comparison between Go uncertain and Go certain before the preparatory stimulus

| Cluster n° | Electrodes | Frequency (Hz) | Time (ms) | Statistic | p-value |
| --- | --- | --- | --- | --- | --- |
| 1 | CPz, CP2, CP4, CPP2h, CPP4h, CPP6h, Pz, P2, P4, PPO1h, PPO2h, PPO6h, POz, PO4 | 7-12 | -500 -  -300 | 334 | 0.004 |

**Supplementary Table 5:** TFR cluster obtained by comparison between Go uncertain and Go certain in reponse to the imperative stimulus

| Cluster n° | Electrodes | Frequency (Hz) | Time (ms) | Statistic | p-value |
| --- | --- | --- | --- | --- | --- |
| 1 | AF3, AFz, AF4, AFF5h, AFF1h, AFF2h, AFF6h, F7, F5, F3, F1, F6, F8, FFT9h, FFT7h, FFC5h, FFC3h, FFC1h, FFT8h, FFT10h, FT7, FC5, FC3, FC1, FC6, FT8, FTT9h, FTT7h, FCC5h, FCC3h, FCC1h, FCC6h, FTT8h, FTT10h, T7, C5, C3, C1, C4, C6, T8, TTP7h, CCP5h, CCP3h, CCP1h, CCP4h, CCP6h, TTP8h, TP7, CP5, CP3, CP1, CPz, CP2, CP4, CP6, TP8, TPP7h, CPP5h, CPP3h, CPP1h, CPP2h, CPP4h, CPP6h, TPP8h, TPP10h, P7, P5, P3, P1, Pz, P2, P4, P6, P8, PPO5h, PPO1h, PPO2h, PPO6h, PO3, POz, PO4, POO2 | 6 - 22 | 250 - 500 | -460 | 0.0001 |
| 2 | AF3, AFz, AF4, AFF5h, AFF1h, AFF2h, AFF6h, F5, F3, F1, Fz, F2, F4, F6, F8, FFC1h, FFC2h, FFC4h, FFC6h, FFT8h, FFT10h, FC1, FCz, FC2, FT8, FCC3h, FCC1h, FCC2h, FTT10h, C3, C1, Cz, C2, CCP5h, CCP3h, CCP1h, CCP2h, CCP4h, CP5, CP3, CP1, CPz, CP2, CP4, CPP5h, CPP3h, CPP1h, CPP2h, CPP4h, P5, P3, P1, Pz, P2, P4, P6, PPO5h, PPO1h, PPO2h, PPO6h, PO7, PO3, POz, PO4, PO8, POO1, POO2, O1, Oz, O2 | 3 - 15 | 0 - 400 | 2901 | 0.0001 |

**Supplementary Table 6:** TFR cluster obtained by comparison between NoGo and Go uncertain in reponse to the imperative stimulus

| Cluster n° | Electrodes | Frequency (Hz) | Time (ms) | Statistic | p-value |
| --- | --- | --- | --- | --- | --- |
| 1 | AF3, AFz, AF4, AFF5h, AFF1h, AFF2h, AFF6h, F7, F5, F3, F1, Fz, F2, F4, F6, F8, FFT9h, FFT7h, FFC5h, FFC3h, FFC1h, FFC2h, FFC4h, FFC6h, FFT8h, FFT10h, FT7, FC5, FC3, FC1, FCz, FC2, FC6, FT8, FTT9h, FTT7h, FCC5h, FCC3h, FCC1h, FCC2h, FCC6h, FTT8h, FTT10h, T7, C5, C3, C1, Cz, C2, C6, T8, CCP5h, CCP3h, CCP1h, CCP2h, CCP6h, TTP8h, CP3, CP1, CPz, CP2, CP4, CP6, TP8, CPP5h, CPP3h, CPP1h, CPP2h, CPP4h, CPP6h, TPP8h, P5, P3, P1, Pz, P2, P4, P6, P8, PPO5h, PPO1h, PPO2h, PPO6h, PO7, PO3, POz, PO4, PO8, POO9h, POO1, POO2, O1, O2 | 4 - 22 | 0 - 450 | 11022 | 0.0001 |
